# Defining the relative and combined contribution of CTCF and CTCFL to genomic regulation

**DOI:** 10.1101/2019.12.14.874560

**Authors:** Mayilaadumveettil Nishana, Caryn Ha, Javier Rodriguez-Hernaez, Ali Ranjbaran, Erica Chio, Elphege P. Nora, Sana B. Badri, Andreas Kloetgen, Benoit G. Bruneau, Aristotelis Tsirigos, Jane A. Skok

## Abstract

**Background:** Ubiquitously expressed CTCF is involved in numerous cellular functions, such as organizing chromatin into TAD structures. In contrast, its paralog, CTCFL is normally only present in testis. However, it is also aberrantly expressed in many cancers. While it is known that shared and unique zinc finger sequences in CTCF and CTCFL enable CTCFL to bind competitively to a subset of CTCF binding sites as well as its own unique locations, the impact of CTCFL on chromosome organization and gene expression has not been comprehensively analyzed in the context of CTCF function. Using an inducible complementation system, we analyze the impact of expressing CTCFL and CTCF-CTCFL chimeric proteins in the presence or absence of endogenous CTCF to clarify the relative and combined contribution of CTCF and CTCFL to chromosome organization and transcription.

**Results:** We demonstrate that the N terminus of CTCF interacts with cohesin which explains the requirement for convergent CTCF binding sites in loop formation. By analyzing CTCF and CTCFL binding in tandem we identify phenotypically distinct sites with respect to motifs, targeting to promoter/intronic intergenic regions and chromatin folding. Finally, we reveal that the N, C and zinc finger terminal domains play unique roles in targeting each paralog to distinct binding sites, to regulate transcription, chromatin looping and insulation.

**Conclusion:** This study clarifies the unique and combined contribution of CTCF and CTCFL to chromosome organization and transcription, with direct implications for understanding how their co-expression deregulates transcription in cancer.

## Introduction

CTCF is involved in numerous cellular functions, some of which can be attributed to its role in organizing chromatin into TAD structures. The latter involves a loop-extrusion mechanism whereby cohesin rings create loops by actively extruding DNA until the complex finds two CTCFs bound in convergent orientation, which block its movement [1-5]. While it is known that convergently orientated CTCF binding sites preferentially form loops while divergent sites delineate boundary regions [6], it is not clear why convergently, rather than divergently orientated CTCF sites can stop the movement of cohesin on chromatin. CTCF can also act as a transcription factor (TF) controlling the expression of many genes by binding to their TSSs [7]. In addition, CTCF can pause transcription. Thus, it is clear that not all CTCF sites are created equal and there are site-specific functional distinctions, but it is not known whether these can be attributed to differences in binding site motifs and/or the action of cofactors that bind CTCF.

CTCFL (CTCF like), otherwise known as BORIS (Brother of the Regulator of Imprinted Sites), is the paralog of CTCF [8]. It emerged by gene duplication of *CTCF* during evolution in the ancestry of amniotes [9]. In contrast to CTCF, which is a constitutively and ubiquitously expressed essential protein, CTCFL is expressed only transiently in pre-meiotic male germ cells of healthy individuals together with CTCF [10]. It plays a unique role in spermatogenesis by regulating expression of pluripotency and testis specific genes [10-12]. It is also aberrantly activated in cancers of several lineages including lung [13-15], breast [16, 17], uterine [18], esophageal [19], hepatocellular [20], ovarian [21-24], prostate [25], urogenital [26] and neuroblastoma [27]. CTCFL has been shown to promote neoplastic transformations by its interference in cellular processes including invasion and apoptosis, cell proliferation and immortalization [21, 22, 27-29]. Furthermore, CTCFL was identified as one of the most promising cancer testis antigens by the NCI [30] and it is known to be important in activating the expression of numerous other cancer testis antigens.

CTCF is bound to chromatin through a subset of its eleven zinc fingers (ZFs). The core ZFs 3-7 make sequence specific contacts with DNA and it is thought that ZFs 8 and 9 provide stability [31, 32]. Together, the 11 zinc fingers of CTCF contribute to its multivalent nature and ability to bind to about 50,000 sites across the genome [33]. The DNA binding ZF region of CTCF and CTCFL share 74% sequence identity [9], however, the N- and C-terminal domains are quite distinct and likely interact with different binding partners that contribute to their unique functions [34]. CTCFL has the ability to bind to and compete with CTCF at a subset of its binding sites, owing to the similarity in the DNA binding region [10, 35]. Although differences in the two proteins can lead to divergent and antagonistic effector functions [10], little is known about the mechanisms underlying these different outcomes.

In this context, it is not clear how the N/C terminals and zinc finger domains contribute to CTCF’s site-specific roles and which regions of the protein are involved in interacting with cohesin. There is contradictory evidence supporting and disputing a role for the C terminal region of CTCF in mediating CTCF-cohesin interaction, respectively from the Felsenfeld and Reinberg labs [36, 37]. Furthermore, the issue of which region of CTCF halts cohesin’s movement on chromatin remains an unsolved problem as co-immunoprecipitation or ChIP-seq analysis of mutants lacking these or other domains has not been published. It is also not known which part/s of CTCFL are important for its role in gene regulation and whether the individual domains have distinct functional impacts at different binding sites. Like CTCF, CTCFL can act as a transcription factor (TF), but given that its binding does not overlap with cohesin [10, 35], it is unlikely to be able to phenocopy CTCF’s function in acting as an insulator at boundary sites, but this has not been analyzed. Pertinent to our investigations is the finding that CTCFL can bind competitively to a subset of sites that CTCF binds [10, 35] and because of the likely differences in the insulating capability of the two proteins, eviction of CTCF at these sites could have an impact on chromosome architecture linked to changes in gene regulation, but this has not been examined.

Investigating the impact of CTCFL overexpression in cancer cells is difficult because of the confounding effects of other genetic and epigenetic alterations. To circumvent these issues, we combined use of a CTCF degron system (which acutely and reversibly depletes endogenous CTCF) [7], with knocked-in doxycycline inducible transgenes encoding intact CTCFL and CTCF- CTCFL chimeric proteins at the *Tigre* locus. This dual system allowed us to elucidate the interplay of CTCFL and CTCF by analyzing the functional impact of each protein in cells where they were expressed individually or together. Using this approach, we highlight an interesting aspect of functional importance: not all CTCF and CTCFL binding sites are created equal. CTCF and CTCFL each bind to a set of unique and overlapping sites that have distinct DNA motifs, chromatin folding properties and biases for being in promoters rather than intronic or intergenic regions. Expressing CTCF-CTCFL chimeric proteins with swapped N and C terminal domains revealed that the zinc finger region of both CTCF and CTCFL defines their respective DNA motif specificity, while the N and C terminal domains influence whether the proteins bind promoters or intergenic and intronic regions. In contrast to the effect on CTCF binding [38], we demonstrate that RNA degradation does not hamper the DNA binding properties of CTCFL. We also show that CTCF and CTCFL have distinct impacts on chromatin folding: while CTCF demarcates TAD boundaries, CTCFL cannot insulate chromosome domains which is explained by its inability to physically interact with cohesin. Finally, we establish that both the zinc fingers and N terminal region of CTCF contribute to insulation at domain boundaries although cohesin only binds to the N terminal region of this protein. The latter finding is consistent with the role that convergently bound CTCF proteins plays in loop formation. This study clarifies the relative and combined contribution of CTCF and CTCFL to chromosome organization and transcription, with direct implications for understanding how their co-expression deregulates transcription in cancer.

## Results

### System to investigate the interplay between CTCFL and CTCF in somatic cells

CTCF and CTCFL have similar zinc finger domains but their N and C terminal regions have no homology as indicated in **Fig. 1a**. In line with these differences CTCF and CTCFL have very different expression profiles: CTCF is present in all cell types, while in contrast CTCFL is normally only expressed transiently in pre-meiotic male germ cells (**Additional File 1: Figure S1A**). However, CTCFL is aberrantly activated in a wide variety of cancer types [8] and publicly available data from genomic studies demonstrates that, in the context of cancer, CTCFL exhibits a variety of genetic alterations. As of July 2019, 382 of the 10950 (3%) cancer samples profiled in cBioPortal [39] were found to have genetic changes in *CTCFL*, with amplification occurring most frequently (58%) in these patient samples (**Additional File 1: Figure S1B-E**). Moreover, there is a clear correlation between amplification of *CTCFL* and its increased expression in several cancer types including ovarian, uterine, cervical, lung squamous and head and neck cancer (**Additional File 1: Figure S1E**).

**Fig. 1.**
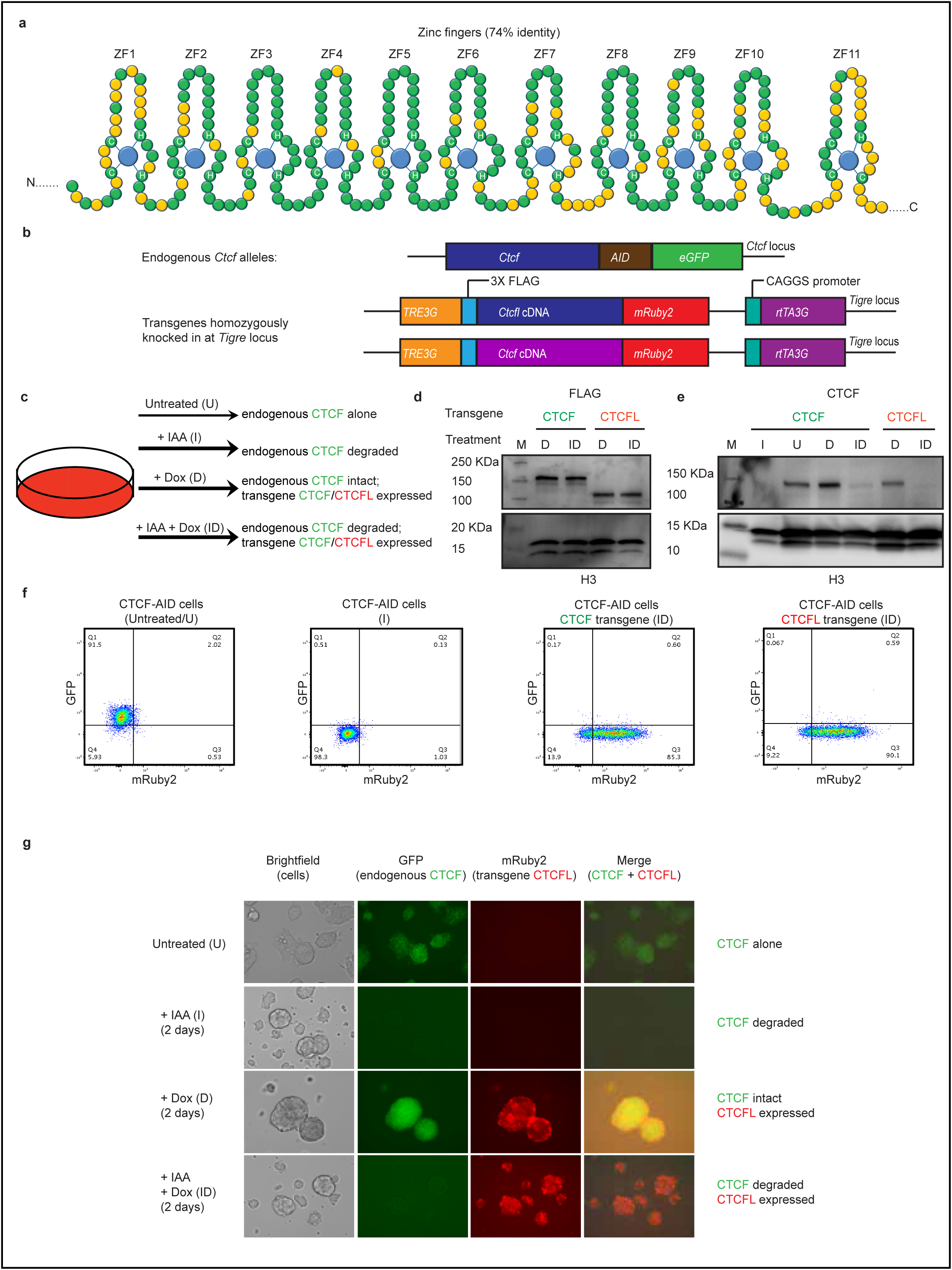
System to investigate the interplay between CTCFL and CTCF in somatic cells. **a** Schematic representation of the similarities and differences between CTCF and CTCFL. Figure adapted from Marshall et al. [34]. The DNA binding domain of both proteins is composed of 11 zinc fingers. ZFs 1-10 and ZF11 belong to the C_2_H_2_ and C_2_HC class of ZFs, respectively. Shared and different amino acids in CTCF and CTCFL are shown in green and yellow, respectively. Blue circles indicate zinc ions and histidines and cysteines that form coordinate bonds with zinc are marked. **b** Scheme of genetic modifications in the *Ctcf* locus and the doxycycline inducible transgenic *Ctcfl* or *Ctcf* knocked-in at the *Tigre* locus. The endogenous *Ctcf* contains the auxin inducible degron (AID) and the eGFP tag on both alleles. Both *Ctcf* and *Ctcfl* transgenes harbor an N terminal 3 X FLAG tag and C terminal mRUBY2 as well as *TetO-3G* element and *rtTA3G* for doxycycline induced expression. **c** Experimental strategy for expression of dox-inducible CTCF/CTCFL transgenes in the presence and absence of CTCF using the auxin inducible degron system. Addition of indole-3- acetic acid (IAA) a chemical analog of auxin leads to transient and reversible degradation of CTCF, while addition of doxycycline (Dox) leads to induction of the respective transgene expression. The four conditions used in our analysis are: U, untreated cells; I, IAA treated for CTCF depletion; D, Dox induced expression of transgenic CTCF/CTCFL; ID, IAA plus Dox treated for depletion of CTCF and induction of transgene expression. **d** Western blot using FLAG antibody shows that the level of expression of transgenes are comparable across the cell types (CTCF and CTCFL in D and ID conditions). CTCF has a predicted molecular weight of 84 KDa and CTCFL, 74 KDa. However, CTCF is known to migrate as a 130 kDa protein [83]. Since the transgenes are expressed as fusion proteins with FLAG tag and mRuby2, which together adds another 29 kDa, the resulting proteins migrate at 159 and 103 KDa respectively. **e** Western blot with CTCF antibody shows the presence of endogenous and transgene CTCF. Histone H3 serves as a loading control in (**d)** and (**e). M** is the molecular weight ladder and the molecular weights are marked. **f**,**g** Flow cytometry and Microscopy confirmedß that the level of mRuby2 expression of transgenic CTCF and CTCFL are comparable.

Despite the finding that CTCFL is aberrantly expressed in numerous cancers, little is known about its impact on chromatin organization and gene regulation and the mechanism underlying its effector functions and interplay with CTCF. To address these questions, we made use of an auxin-inducible degron (AID) mESC system in which we could study the effects of CTCFL in the presence and absence of CTCF [7]. In this system both endogenous CTCF alleles are tagged with AID as well as eGFP (CTCF-AID-eGFP) (**Fig. 1b**) and they constitutively express the auxin-activated ubiquitin ligase TIR1 (from *Oryza sativa*) from the *Rosa26* locus. Addition of indole acetic acid (IAA; an analogue of auxin) leads to rapid poly-ubiquitination and proteasomal degradation of the proteins tagged with the AID domain. Degradation after 48 h of auxin treatment in treated and control cells was confirmed by Western blot, fluorescence microscopy and flow cytometry (**Fig. 1c-g**). To study the impact of CTCFL expression, we established a rescue system wherein the ESC degron cell line was modified to individually express either a stable doxycycline-inducible *Ctcfl* or control wild-type *Ctcf* transgene from the *Tigre* locus (**Fig. 1b, c**). Individual clones with comparable transgenic expression levels were selected based on Western blot and FACS analysis (**Fig. 1d-f**). The four conditions used for our analysis are shown in (**Fig. 1c and g**).

### Distinct characteristics of CTCF and CTCFL and their binding sites

While it is known that CTCF and CTCFL bind to both unique and overlapping binding sites [10, 35], it is not known whether these sites have distinct properties and whether binding of the two proteins at different locations leads to distinct effector functions. Furthermore, it is not known whether the presence versus absence of CTCF alters the profile of CTCFL binding and / or its impact on gene expression. Use of the dual CTCF degron system combined with expression of transgenic CTCF or CTCFL provided us with a unique system with which to address these questions. We first performed ChIP-seq (by ChIPmentation) to examine how DNA binding of transgenic CTCF or CTCFL changes in the presence (D) and absence (ID) of endogenous CTCF, using the FLAG tag in the transgenes. We also performed RNA-seq.

RNA seq and ChIP-seq confirmed that transgenic CTCFL/CTCF expression and binding occur only after doxycycline induction (**Fig. 2a**). As expected, we found locations where CTCFL bound to unique sites and sites where it overlapped with CTCF binding. Binding of CTCFL was detected at the promoter and an intragenic site in *Ctcfl*. CTCF was absent at this site but was bound to an intragenic site overlapping CTCFL binding, as well as two other CTCF sites within the *Ctcfl* gene. ChIP-seq and RNA-seq indicated that binding of CTCFL at promoters of genes including testis specific *Stra8 and Prss50* was linked to their activation. STRA8 and CTCFL have an overlapping expression pattern while PRSS50 is expressed during spermatogenesis in CTCFL positive cells as well as in subsequent stages of development when CTCFL is no longer expressed [10] (**Fig. 2a**). Binding within exons of some genes such as *Gal3st1* (**Additional File 1: Figure S2A**) was correlated with an increase in transcriptional output [10, 40], while binding at promoters of other genes (*Rapgef1*) (**Additional File 1: Figure S2A**) was not. Thus, CTCFL binding does not always impact gene expression. Other loci (e.g. the *Hoxb cluster*) exhibited a preference for CTCF rather than CTCFL binding (**Additional File 1: Figure S2B**).

**Fig. 2.**
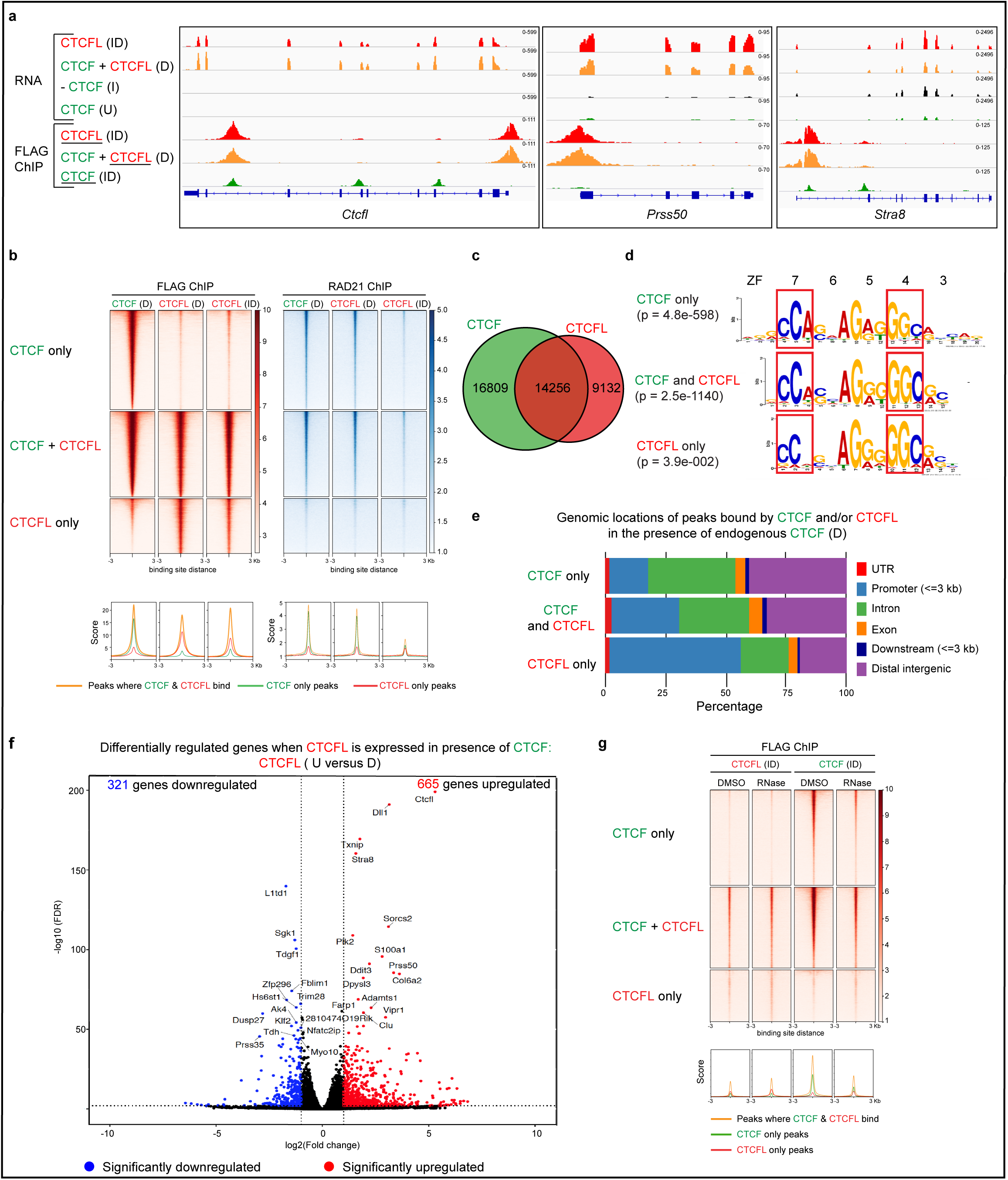
Distinct characteristics of CTCF and CTCFL and their binding sites. **a** IgV tracks show RNA-seq in cells harboring the CTCFL transgene in U, I, D and ID conditions. FLAG ChIP-seq to detect CTCFL and CTCF in cells with the respective transgenic knock-ins. The protein whose binding is being assessed is underlined. Expression at *Ctcfl, Prss50* and *Stra8* are also shown. **b** Heatmaps showing CTCF and CTCFL ChIP-seq signals at regions where they bind alone or together in CTCF D, CTCFL D and ID conditions. The heatmaps are divided into CTCF only, CTCF + CTCFL overlapping and CTCFL only sites. Cohesin binding profiles (RAD21 ChIP) are shown for the corresponding conditions. Peaks are ranked by FLAG ChIP in cells expressing the CTCF transgene. Average profiles are shown below the corresponding heatmaps. **c** Venn diagrams showing the distribution of unique or overlapping CTCF and CTCFL binding sites (D condition). **d** Binding site motifs for unique or overlapping CTCF and CTCFL binding sites. Zinc fingers and the corresponding bases to which they bind are marked. **e** Annotation of the genomic locations of peaks for unique or overlapping CTCF and CTCFL binding sites in D condition. The locations of UTR, promoters (+/- 3 kb around TSS), introns, exons, downstream (3 kb) and distal intergenic regions are marked. **f** Volcano plot highlighting DEGs in wild-type versus CTCFL expressing mESCs in the presence of endogenous CTCF. Red and blue points identify genes with significantly increased or decreased expression, respectively (FDR<0.01). The nu mber of genes that are significantly up or downregulated is indicated in either case. **g** Heatmaps showing the inhibition of chromatin binding of CTCF, but not CTCFL, by RNaseA treatment. The heatmaps are divided into CTCF only, CTCF + CTCFL overlapping and CTCFL only sites. Peaks are ranked by FLAG ChIP in cells expressing the CTCF transgene (CTCF-ID). Average profiles are shown below the corresponding heatmaps.

To further examine the unique and overlapping binding sites of CTCF and CTCFL we performed FLAG ChIP-seq in the presence of endogenous CTCF (CTCFL D condition) (**Fig. 2b, c**). We identified 16,809 CTCF-only and 9,132 CTCFL-only binding sites, while both proteins shared 14,256 sites. In total, 46% of CTCF bound sites were occupied by CTCFL and 61% of CTCFL bound sites were bound by CTCF (**Fig. 2c**). In agreement with previous studies [10, 35, 41-43], we found that CTCF and cohesin have an overlapping binding profile as CTCF blocks the cohesin mediated extrusion of DNA **(Fig. 2b**). RAD21 was localized only at sites where CTCF normally binds i.e. CTCF only and CTCF + CTCFL sites, but it failed to localize at CTCFL only sites (**Fig. 2b**). Compared to untreated cells, induction of CTCFL (CTCFL D) did not lead to a drastic alteration in the RAD21 profile. However, when CTCFL was expressed in the absence of CTCF (CTCFL ID), RAD21 peaks were globally depleted. As a result, it is unlikely that CTCFL will have the ability to phenocopy CTCF’s function as insulator.

Motif analysis revealed that CTCF only sites contained the consensus CTCF motif (JASPAR MA0139.1). CTCFL only sites had less of a requirement for the ‘A’ base in the triplet where ZF7 binds as shown previously [10], as well as an increase for ‘C’ in the triplet that was bound by ZF4 (**Fig. 2d**). The change in the ZF7 binding region mirrors differences between ZF7 in CTCF and CTCFL (**Fig. 2d, 1a**). Changes in the dependence of ‘C’ in the triplet where ZF4 binds could either be explained by differences in ZF4 between CTCF and CTCFL and/or by changes in binding of ZF7 affecting upstream binding of ZF4. Overlapping CTCF and CTCFL binding sites had a motif intermediate to that of CTCF and CTCFL only sites.

In order to define the functional significance of the three different bindings sites (CTCF only, CTCF + CTCFL overlapping and CTCFL only), we compared their genomic distribution. CTCF only sites have a preference for intronic and intergenic regions, while in contrast, CTCFL-only sites favored promoters (**Fig. 2e**). Overall, only 22% of CTCF sites were at promoters, versus 38% for CTCFL (**Fig. 2e**). CTCF binding at promoters co-occurred with CTCFL in 60% of cases, while CTCFL binding at promoters occurred without CTCF in 55% of cases.

When CTCFL was expressed in the presence of endogenous CTCF, a total of 986 genes were significantly deregulated (lfc > 1 and fdr < 0.01) (**Fig. 2f, Additional File 2: Table S1**). Most CTCFL-regulated genes are not controlled by CTCF and vice versa, since there was little overlap between genes deregulated by induction of CTCFL in the absence of endogenous CTCF (CTCFL U versus ID) and those altered by depletion of endogenous CTCF (CTCF U versus I) (**Additional File 1: Figure S2C-E**). Of interest, 146 out of 219 genes found in the overlapping subset were regulated by CTCF and/or CTCFL binding at the promoters, 76 of which had overlapping binding sites. Analysis of CTCF (U versus I) and CTCFL (U versus D) mediated changes in gene expression in the context of promoter binding demonstrated 36.2% and 50.8% of alterations, respectively (**Additional File 1: Figure S3A**). Because CTCF was bound to many more sites than CTCFL (**Fig. 2c**) there was an increase in the number of genes deregulated in the CTCFL (U versus ID) cohort compared to the CTCFL (U versus D) subset (**Fig. 2f, Additional File 1: Figure S2C**,**D**). It is important to note that without the degron system it would not have been possible to examine the interplay between CTCF and CTCFL.

Taken together these data demonstrate that CTCFL has more of a preference for binding promoters than CTCF, which highlights the functional differences of the two factors. Furthermore, when CTCF is located at promoters, we found that binding preferentially occurs at CTCF + CTCFL overlapping sites, suggesting that CTCFL binding sites may be functionally distinct from the CTCF only bound sites.

### Interaction with RNA is not essential for the binding of CTCFL to chromatin

A recent study has shown that interaction with RNA is essential for binding of CTCF to DNA [38]. The zinc fingers of CTCF have two RNA Binding Regions (RBRs) that facilitate RNA interaction. One RBR extends from amino acids 264-275 that stretch from nearly the end of the N terminal domain through ZF1 and the other encompasses amino acids 536-544 in ZF10 [36, 38]. It is of note that there are considerable differences between the sequences of ZF1, ZF10 and ZF11 in CTCF versus CTCFL. The RBR at ZF1 (KTFQCELCSYTCPR) of CTCF shows a clear difference in sequence from that of CTCFL (**G**TF**H**C**DV**C**MF**T**SS**R, differences bolded), while the RBR at ZF10 (QLLDMHFKR) is relatively conserved (QLL**NA**HF**RK**). Deletions of both RBRs were shown to disrupt DNA binding, with the mutation of ZF10 having less of an impact than that of ZF1 [38]. In addition, the C terminal 576-611 amino acids that connect the C terminal domain of CTCF with ZF 11 (also an RBR), have been shown to be important for the diffusion, clustering, target search and self-association of CTCF. This RBR region does not physically interact with cohesin, but contributes to the formation of CTCF clusters in an RNA dependent manner and these clusters block extruding cohesin [54, 55]. Since the RBR’s are not significantly conserved between CTCF and CTCFL, we sought to determine if RNA has any role to play in the binding of CTCFL to DNA by treating cells expressing transgenic CTCF (CTCF-ID) and CTCFL (CTCFL-ID) with RNase. FLAG ChIP-seq revealed that as expected, CTCF exhibited reduced binding to chromatin [38], while in contrast, CTCFL binding was unaltered **(Fig. 2g)**. Thus, CTCFL does not require RNA to bind chromatin.

### CTCFL activates cancer testes antigens (CTA) and components of cancer relevant signaling pathways

CTCFL, itself a cancer testis antigen (CTA) referred to as Cancer/Testis Antigen 27 has an impact on expression of other CTA genes. Indeed, upon induction with Dox, *Ctcfl* and *Dll1* were the most highly upregulated genes (**Fig. 2f**). DLL1 is a Notch ligand known to play a major role in cancers like breast cancer [44, 45] and squamous neoplasias [46] and it is thought to be a promising therapeutic target [47]. Our ChIP-seq data showed CTCFL binding at the promoter of *Dll1* and other CTAs such as *TSP50 or Prss50* [48] and those belonging to the MAGE family (MAGE-B4, MAGE- E1, MAGE-F1) (**Additional File 1: Figure S3A, B**). CTCFL binding was also linked to increased expression of the ADAM family of proteins (ADAMTS2, ADAMTS15) (**Additional File 1: Figure S3C**). Use of the degron system allowed us to determine whether there is overlap in the genes that CTCF and CTCFL regulate, e.g. *Stra8* **(Fig. 2a**), or whether control is mutually exclusive, as in the case of the other genes highlighted above (**Additional File 1: Figure S3A-C**).

Previous studies have shown that CTCFL transgenic mice die within a few hours after birth. They exhibit ocular hemorrhaging and unfused eyelids, a phenotype typical of mouse models in which the TGFβ pathway is deregulated. In line with this, RNA-seq analysis of ES cells from the mice revealed upregulation of TGFβ1 [40]. Our ChIP-seq and RNA-seq data demonstrated that CTCFL binds to the promoter of *Tgfβ1*, leading to its upregulation (**Additional File 1: Figure S3D**). These findings highlight the links between CTCFL and TGFβ1. Additionally, a subset of TGFβ1 target genes (*Bhlhe40, Klf10, Gadd45b*) were upregulated in our dataset [49-51] (**Additional File 1: Figure S3E**). Furthermore, *Stat1*, a protein with both tumor suppressor and oncogenic properties [52] was bound and activated by CTCFL (**Additional File 1: Figure S3D**). We also identified upregulation of *Cited1*, which encodes Cbp/p300-interacting transactivator 1 that is a cofactor of the p300/CBP-mediated transcription complex [53] (**Additional File 1: Figure S3F**). These data demonstrate that ectopic expression of CTCFL is sufficient to trigger expression of a panel of genes that regulate several signaling pathways important in cancer.

### The impact of CTCFL on 3D chromatin organization

To determine if CTCFL either shares or antagonizes the role of CTCF in chromosome folding we performed Hi-C (see **Additional File 1: Figure S4A and Additional File 3: Table S2** for quality control (QC) analysis). Consistent with the findings from previous studies our Principal Component Analysis (PCA) showed that compartments, which separate active euchromatin (A compartment) from inactive heterochromatin (B compartment) remain largely unchanged when CTCF was degraded (CTCFL I) [7]. We also could not detect any changes in compartments after induction of CTCFL, either in the presence (CTCFL D) or absence of endogenous CTCF (CTCFL ID) (**Fig. 3a**). Next we examined the impact of CTCFL on the highly self-interacting topologically associated domain (TAD) structures that form independently of compartments. Transgenic expression of CTCFL (CTCFL ID) did not rescue TAD structures that were lost upon CTCF depletion [7]. In contrast, control experiments using cells that harbor the CTCF transgene CTCF (CTCF ID) were able to restore these structures (**Fig. 3b,c**). Consistent with the dose dependent effects of CTCF [7] we found that TADs were strengthened by expressing the CTCF transgene in the presence of endogenous CTCF (CTCF D) while expression of CTCFL in the presence of endogenous CTCF (CTCFL D) did not dramatically alter TAD structure at a global level (**Fig. 3b,c**).

**Fig. 3.**
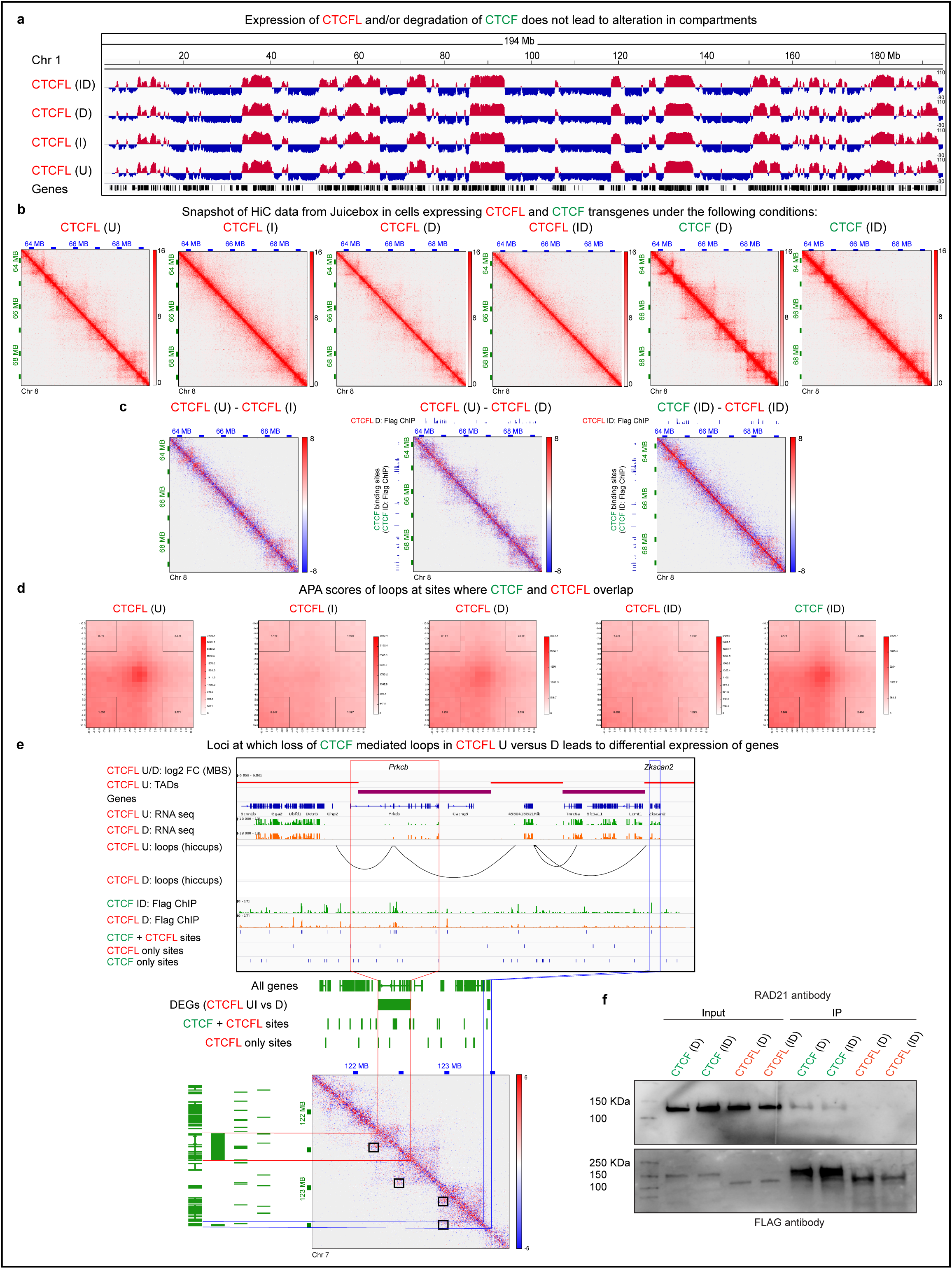
The impact of CTCFL on 3D chromatin organization. **a** IgV tracks showing principal component analysis characterizing the A/B status of compartments (Red track: A compartment, PC1>0; Blue track: B compartment, PC1<0) in cells harboring the CTCFL transgene under U, I, D, and ID conditions. Data from chromosome 1 is shown. **b** Hi-C data from Juicebox corresponding to Chr 8: 63,616,214-69,456,200 8: 63,566,214-69,406,200 at 10 kb resolution. TADs show up as triangles on Hi-C contact maps whose intensity represents interaction strength. Heatmap of Hi-C interactions demonstrates loss of TADs following CTCF depletion (CTCFL I). CTCFL expression does not have a major impact on global TAD structure in the presence (D) or absence (ID) of CTCF. Strengthening of TADs is seen in CTCF D and rescue of TADs in CTCF ID. **c** Subtraction heatmaps of Hi-C data from Juicebox corresponding to CTCFL (U – I), CTCFL (U - D) and CTCF ID – CTCFL ID. CTCF binding sites (CTCF ID: FLAG ChIP) are shown on the y-axis and FLAG ChIPs of CTCFL-D and -ID on the x-axis as indicated. **d** Aggregate Peak Analysis demonstrates the strength of the loops at sites where CTCF and CTCFL bind competitively. The transgenes and the respective treatments are indicated. The color intensity at the center of the plot is indicative of loop strength. APA scores are shown in the corners. Values > 1 indicate presence of loops. Examples of altered loops at specific loci are shown in **Additional File 1: Figure S4B. e** Screenshots of UCSC genome browser showing features of chromatin organization including Mean Boundary Scores (MBS), presence of TADs and alterations in loops as well as RNA and ChIP-seq tracks. The ChIP- seq peaks unique to CTCF and CTCFL as well as those that are overlapping are shown. The transgenes harbored by the cells and the respective treatments are indicated. Lower panel shows a representative case where alterations in loops and differential expression of genes occur at sites where CTCF and CTCFL binding overlaps. A snapshot of subtraction heatmap from Juicebox is shown with the loops highlighted in boxes. Loops appear as dots at the apex of TADs, the intensity of which defines the ‘loop strength. **f** Co-IP experiments showing the interaction of RAD21 with transgenic CTCF and CTCFL in both D and ID conditions.

Since CTCFL does not bind everywhere that CTCF binds, we focused on overlapping binding sites and performed an aggregate peak analysis (APA) to estimate the strength of the loops at these locations [56]. In this evaluation, the signals from a set of peak pixels are superimposed such that the color intensity corresponds to the strength of the loops. Cells with intact CTCF (CTCF U) had the strongest loops and as expected, these disappeared when CTCF was degraded (CTCF I) and could be rescued by expression of control transgenic CTCF (CTCF ID). In contrast, CTCFL was unable to rescue CTCF-mediated looping (CTCFL ID) (**Fig. 3d**). An example of this is shown in **Additional File 1: Figure S4B**. Furthermore, expression of CTCFL in the presence of CTCF, reduced loop strength, indicating that binding of CTCFL at CTCF overlapping sites impairs loop formation (**Fig. 3d**). This demonstrates that CTCFL does not have the same function as CTCF in chromosome organization and ectopic CTCFL expression disrupts CTCF-mediated genome folding.

We next asked if the changes in loop strength that were seen at sites where CTCF and CTCFL binding overlaps had any functional impact on transcriptional output. In untreated cells (U), the region corresponding to the *Prkcβ* gene is involved in two loops, one of which has CTCF-CTCFL overlapping binding sites at both anchors and the other only at one of the anchors. Induction of CTCFL led to the disappearance of the loops as well as concomitant overexpression of *Prkcb* (**Fig. 3e**). Upregulation of PRKCβ is of interest because it is a protein implicated in several cancers including lymphoma, glioblastoma, breast, prostate and colorectal cancers [57]. In the same snapshot, downregulation of *Zkscan2* (that encodes a zinc finger with KRAB and SCAN domain protein) is linked to loss of a loop that has a CTCF-CTCFL overlapping binding site at an anchor adjacent to the gene (**Fig. 3e**).

In sum, these analyses demonstrate that CTCFL cannot rescue TAD structure and loop strength that are lost after CTCF depletion. Furthermore, while CTCFL does not have a global impact on TAD structure in the presence of CTCF, it does have an impact on looping at CTCF + CTCFL overlapping sites. Importantly, binding of CTCFL at CTCF + CTCFL overlapping binding sites was linked to differential expression of genes within altered loops. These findings have implications for the role of CTCFL in altering chromatin organization and gene expression in the context of cancer where CTCFL is expressed alongside CTCF.

### CTCFL does not physically interact with cohesin

There is some controversy about which region of CTCF interacts with cohesin. While one report demonstrates physical interaction between the C terminal region of CTCF (amino acids 575 to 611) and the SA2 subunit of cohesin [37], other studies that deleted these amino acids (577-614) showed that they are dispensable [36, 38, 55]. Although RAD21 overlaps with CTCF binding it does not occupy sites bound exclusively by CTCFL (**Fig. 2b**) [10, 35]. It is thus likely that CTCFL fails to physically interact with cohesin, but this has not been directly demonstrated. To investigate, we performed co-immunoprecipitation experiments with lysates from cells induced to express transgenic CTCFL or control transgenic CTCF, in the presence or absence of endogenous CTCF (CTCF D and ID; CTCFL D and ID). We used an antibody to FLAG to pull down transgenic proteins followed by Western blotting with a RAD21 antibody, to determine whether the two proteins interact with the cohesin complex under the different culture conditions. As shown in **Fig. 3f**, we find that RAD21 interacts with CTCF but fails to interact with CTCFL. Surprisingly, we were unable to visualize a RAD21 band in cells induced to express CTCFL in the presence of endogenous CTCF (CTCFL D). This suggests that CTCF and CTCFL may not interact with each other, in contradiction to findings from a previous study [35].

### The role of CTCF and CTCFL zinc fingers and N/C terminal regions in site specific binding

While it is known that the zinc fingers 6 and 7 of CTCF and CTCFL can define site specific selectivity [10, 58], little is known about whether there are other functional contributions made by the zinc fingers or N/C terminal regions of each factor. In order to investigate, we inserted transgenic CTCFL and CTCF with swapped N and C terminal domains into the *Tigre* locus. The fusion proteins (**C**TCF N terminus - CTCF**L** zinc fingers - **C**TCF C-terminus; CTCF**L** N terminus - **C**TCF zinc fingers - CTCF**L** C-terminus) are abbreviated as CLC and LCL where C stands for CTCF and L stands for CTCFL (**Fig. 4a**). The fusion protein transgenes, along with their FLAG and mRUBY tags, were expressed at the same levels as intact transgenic CTCF and CTCFL, as demonstrated by both flow cytometry (**Additional File 1: Figure S5A**) and western blotting (**Fig. 4b**). RNA-seq analysis revealed that each transgenic construct expressed the appropriate domains of CTCFL/CTCF as indicated by peaks at the respective exons (**Additional File 1: Figure S5B**).

**Fig. 4.**
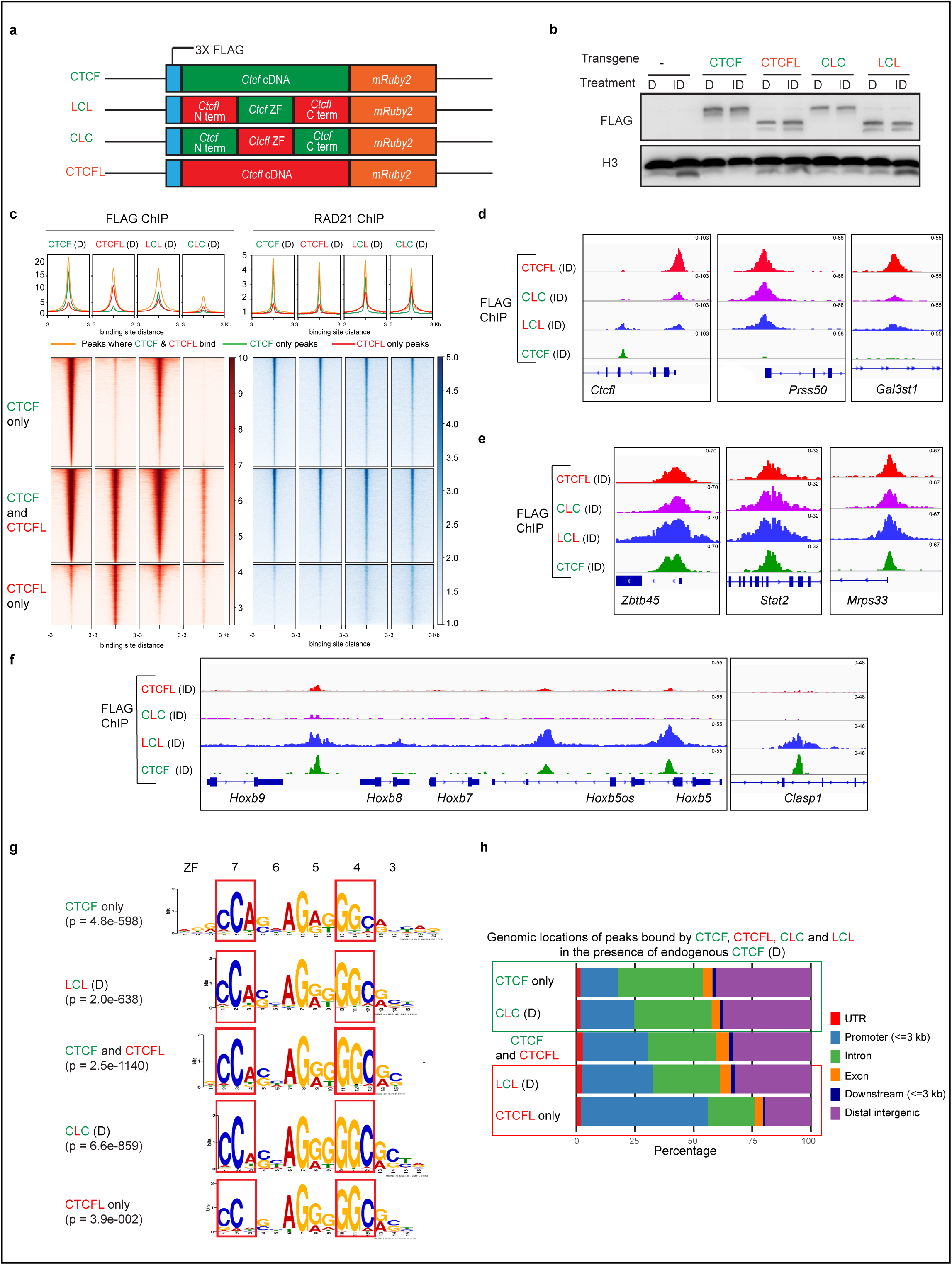
The role of CTCF and CTCFL zinc fingers and N/C terminals in site specific binding. **a** Schematic showing the doxycycline inducible parent (*Ctcf, Ctcfl*) and chimeric (CTCFL N terminus - CTCF ZFs - CTCFL C-terminus (LCL) and CTCF N terminus - CTCFL ZFs - CTCF C- terminus (CLC) transgenes knocked into the *Tigre* locus. All transgenes contain an N terminal 3 X FLAG, C terminal mRUBY2 as well as *TetO-3G* and *rtTA3G* for doxycycline induced expression. **b** Western blot using FLAG antibody shows that the level of expression of transgenes is comparable across cell types as well as experimental conditions (D versus ID). Histone H3 serves as loading control. **c** Heatmaps of CTCF, CTCFL, LCL and CLC ChIP-seq signals at regions where they bind in the presence of endogenous CTCF (D). The heatmaps are divided into CTCF only, CTCF + CTCFL overlapping and CTCFL only regions. Cohesin peaks (RAD21 ChIP) are shown for the corresponding conditions. Peaks are ranked by FLAG ChIP using cells expressing the CTCF transgene. Average profiles are shown above the corresponding heatmaps. **d-f** IgV tracks showing ChIP-seq for CTCF, CTCFL, LCL and CLC at *Ctcfl, Prss50, Gal3st1* (**d**) *Zbtb45, Stat2, Mrps33* (**e**) *Hoxb* and *Clasp1* (**f**) using a FLAG antibody in cells harboring the respective transgenes in the absence of CTCF. **g** Binding site motifs at CTCF and CTCFL only sites, as well as CTCF and CTCFL overlapping sites. Motifs for CLC and LCL were identified in the presence of endogenous CTCF. The zinc fingers and the corresponding bases to which they bind are marked. **h** Annotation of the genomic locations of peaks bound at CTCF and CTCFL only sites, as well as CTCF and CTCFL overlapping sites. Motifs for CLC and LCL were identified in the presence of endogenous CTCF. The locations of UTR, promoter (+/- 3 kb around TSS), intron, exon, downstream (3 kb) and distal intergenic regions are marked.

FLAG ChIP-seq in the presence of endogenous CTCF (D condition) revealed that LCL has a similar binding profile to CTCF, including at CTCF-only sites (**Fig. 4c** and **Additional File 1: Figure S5C, D**). The CTCF zinc fingers are therefore the main determinant of CTCF binding specificity. In contrast, CLC only bound CTCF+CTCFL sites, and was unable to target CTCFL-only sites. CLC also exhibits overall reduced binding compared to CTCFL (**Fig. 4c**). We also found that LCL can bind to a subset of CTCFL only sites where CTCF does not bind (**Additional File 1: Figure S5D**), indicating that the N/C domains of CTCFL participate in targeting to CTCFL-only sites. Interestingly, clustering analysis of binding at CTCF and CTCFL only sites revealed that CLC is able to bind weakly to some CTCF only sites when CTCF is present but not when it is absent (**Additional File 1: Figure S5D**), indicating that the N and C terminals of CTCF might facilitate interaction between CTCF and CLC which in turn could direct binding of CLC to these sites. To test this idea, we performed immunoprecipitation with GFP-Trap magnetic beads in which the endogenous CTCF was pulled down with a GFP antibody and blotted with a FLAG antibody against the transgenes. Our results show a physical interaction of CTCF with itself, CLC and LCL but not CTCFL. CTCF and CLC interaction could explain the presence of CLC at CTCF only sites in the Dox condition (**Additional File 1: Figure S5E**). Examples of chimeric and parent protein binding are shown in the screenshots in **Fig. 4d-f**. CTCFL and CTCF-CTCFL overlapping binding sites are frequently bound by both CLC and LCL (**Fig. 4d, e**), but at some locations fusion protein peaks are reduced in size compared to that of CTCFL (eg. at *Ctcfl, Prss50, Gal3st1* loci) (**Fig. 4d**). Sites bound by CTCF only were preferentially bound by LCL as opposed to CLC (**Fig. 4f**). Interestingly, RAD21 ChIP-seq reveals that CLC and LCL can redistribute cohesin to CTCFL only sites where it does not normally go (**Fig. 4c**). This suggests that both the N/C domain of CTCF (present in CLC) as well as the zinc fingers (present in LCL) participate in how CTCF recruits cohesin.

From the ChIP-seq data we demonstrate that the motif for sites where LCL and CLC bind is similar to that of CTCF and CTCFL, respectively (**Fig. 4g**). These findings indicate that as expected, zinc fingers direct sequence specific binding. In contrast, when we analyzed the genomic annotation intervals (UTR, promoters, introns, exons, downstream and distal intergenic regions) of the fusion protein binding sites, we identified a preference for LCL to be at promoters and CLC to be at intergenic and intronic regions. Thus, LCL resembles CTCFL and CLC resembles CTCF in this aspect of their behavior (**Fig. 4h**). These data reveal that the N and C terminal regions of CTCF and CTCFL contribute functionally to where these factors bind.

Taken together, these data indicate that both the zinc fingers and N and C terminal regions play distinct roles in site directed binding. LCL and CLC resemble CTCF and CTCFL, respectively in terms of their binding motifs highlighting the importance of the zinc fingers. The opposite is the case when it comes to the regions they prefer to bind (promoters versus intergenic and intronic regions): CLC and LCL resemble CTCF and CTCFL, respectively. We further demonstrate that CTCF can interact with itself, CLC and LCL but not CTCFL. Thus, N/C terminals as well as zinc fingers can potentiate dimerization underscoring the functional contributions of each region in this aspect of CTCF biology.

### Gene expression changes of chimeric proteins do not phenocopy that of either parent protein

To determine whether the N and C terminal regions of the CTCF and CTCFL proteins influence transcriptional output we performed RNA-seq on cells expressing LCL and CLC in the presence and absence of CTCF. Fewer genes were deregulated upon induction of LCL (265 genes) and CLC (254 genes) (**Fig. 5a-c, Additional File 2: Table S1**) compared to induction of CTCFL (986 genes) in the presence of CTCF (**Fig. 2f, Additional File 2: Table S1**). Thus, neither CLC nor LCL can phenocopy the impact of CTCFL, underscoring the functional importance of both the zinc finger and N/C terminal domains of this factor. Induction of CLC and LCL in the absence of CTCF, led to an increase in the number of genes that were up and downregulated in each case (**Fig. 5d-f**). However, fewer genes were deregulated than in cells where CTCF was depleted alone (**Additional File 1: Figure S2D**) suggesting that both factors were able to perform a partial rescue.

**Fig. 5.**
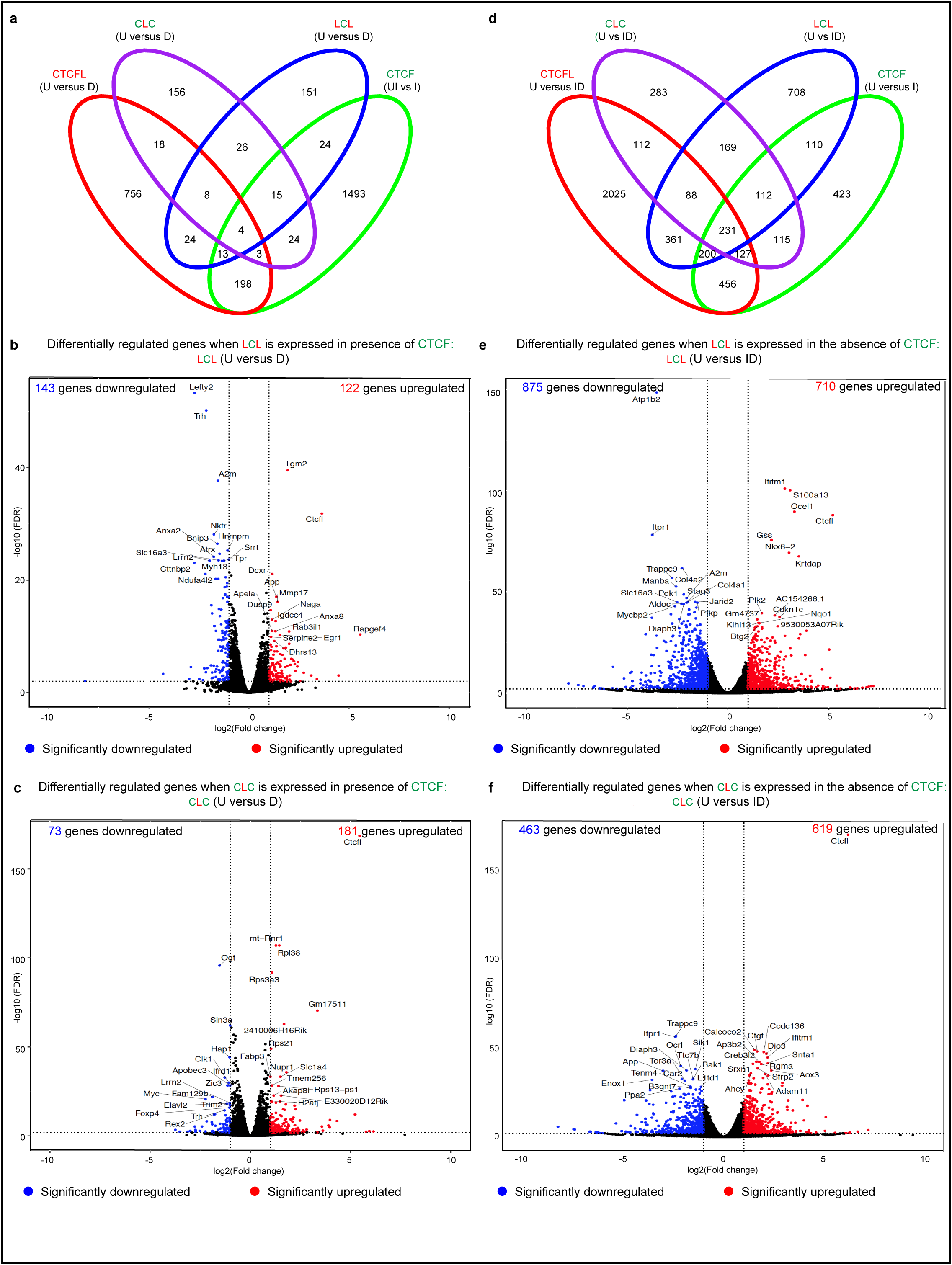
Gene expression changes of fusion proteins do not phenocopy that of either parent protein. **a, d** Venn diagrams showing comparison of deregulated gene expression by CTCFL, LCL and CLC in the presence (**a**) and absence (**d**) of endogenous CTCF with depletion of CTCF. **b, c, e, f** Volcano plot representation of differentially expressed genes in untreated (U) versus LCL (**b, e**) and CLC (**c, f**) expressing mESCs in the presence (D) (**b, c**) and absence (ID) (**e, f**) of endogenous CTCF. Red and blue mark the genes with significantly increased or decreased expression respectively (FDR<0.01). The x-axis shows the log2 fold-changes in expression and the y-axis the log 10 (False discovery rate) of a gene being differentially expressed. The number of genes that are significantly up or downregulated are indicated in either case.

Although the two fusion proteins are largely incapable of phenocopying the impact of the parent proteins on gene expression, there are examples of loci where we see concordant and discordant changes (**Additional File 1: Figure S6**). At *Gadd45g*, concordant changes in gene expression are mediated by CTCFL and CLC suggesting that the zinc finger region of CTCFL is important for the regulation of this gene (**Additional File 1: Figure S6A**). Both parent and fusion proteins are bound upstream of *Gadd45g*, but CTCF removal and induction of LCL have no effect on this gene’s expression status, underscoring the fact that binding does not always equate with proximal changes in gene expression. At the *Igf2* locus, induction of LCL and removal of CTCF leads to its upregulation, indicating that the two factors act discordantly: LCL activates and CTCF represses expression of this gene (**Additional File 1: Figure S6B**). LCL and CTCF bind at an overlapping site upstream of *Igf2os* suggesting this may be a direct effect regulated by the zinc fingers of each protein. Interestingly both CLC and CTCFL appear to activate *Igf2*, although neither factor binds to the upstream region, suggesting an indirect or long-distance effect that results from binding at a distal site. At *Prss50* and *Steap1*, LCL and CLC mediated-activation are concordant with the effects of CTCFL. Furthermore, degradation of CTCF downregulates *Steap1* expression indicating that CTCF is also important for its activation. Here both N/C terminal regions and zinc fingers of CTCFL contribute to the regulation of these genes (**Additional File 1: Figure S6C**). At the *Egr1* locus CLC and LCL act independently of CTCF and CTCFL in regulating transcription (**Additional File 1: Figure S6D**). These examples highlight the fact that overlapping changes in chimeric protein-mediated deregulated genes are not necessarily concordant with expression changes mediated by either parent protein.

### The impact of fusion proteins on chromatin organization

Our finding that CTCFL is unable to rescue the impact of CTCF depletion on TAD structure due to its inability to bind cohesin (**Fig. 3**), begs the question of whether the zinc fingers or N/C terminal domains contribute to this aspect of CTCF’s function. To determine this, we performed Hi-C (see **Additional File 3: Table S2** for QC) and asked if either fusion protein (CLC or LCL) could restore chromatin folding (ID condition). At a global level we observed that transgenic CLC and LCL were partially able to restore the TAD structure that is lost after CTCF depletion (**Fig. 6a, b**). However, CLC and LCL did not have a major impact on TAD structure when expressed in the presence of endogenous CTCF (D condition) (**Additional File 1: Figure S7A**). Additionally, expression of transgenic CTCFL, CLC and LCL in the presence of endogenous CTCF had no significant effect on TAD number or length (**Additional File 1: Figure S7B, C**). As expected we detected fewer, larger TADs upon CTCF depletion and while complete rescue was achieved by expression of transgenic CTCF, no rescue was achieved by induction of CTCFL or the chimeric proteins (**Additional File 1: Figure S7B, C**).

**Fig. 6.**
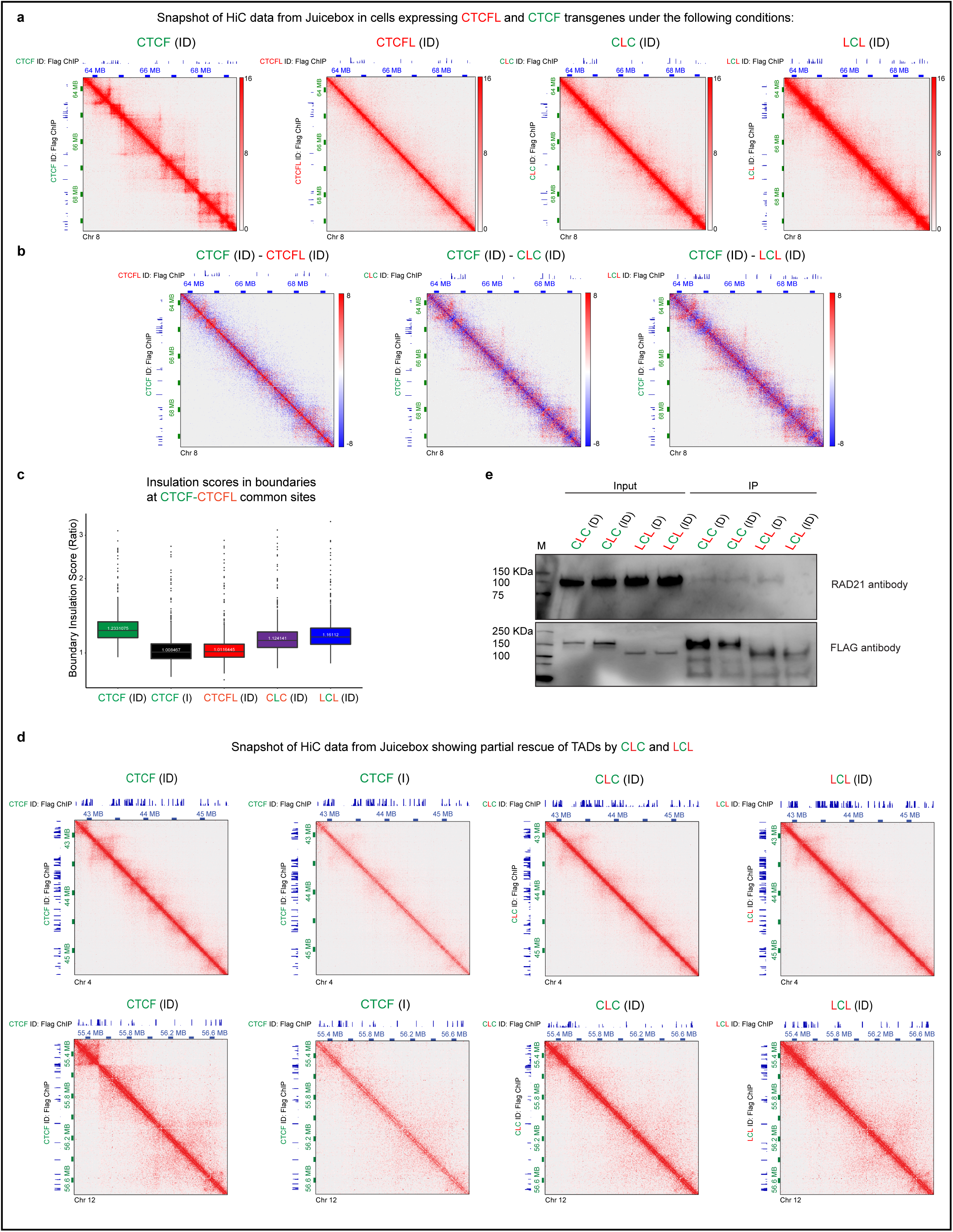
The impact of fusion proteins on chromatin organization. **a** Snapshot of Hi-C data from Juicebox corresponding to Chr 8: 63,616,214-69,456,200 8: 63,566,214- 69,406,200 at 10 kb resolution. Cells harboring CTCF, CTCFL, CLC and LCL transgenes were treated with ID as indicated for 4 days. The corresponding FLAG ChIPs are shown on the x-and y-axis. **b** Subtraction heatmaps of Hi-C data from Juicebox corresponding to CTCF-ID – CTCFL-ID, CTCF-ID – CLC-ID and CTCF-ID – LCL-ID. CTCF binding sites (CTCF-ID: Flag ChIP) are shown on the y-axis and Flag ChIPs of CTCFL-ID, CLC-ID and LCL-ID on the x-axis as indicated. **c** Insulation scores in boundaries of CTCF + CTCFL overlapping sites in CTCF depleted cells (CTCF-I) as well as cells depleted of CTCF but induced to express transgenic CTCF, CTCFL, CLC or LCL (ID condition). **d** Snapshot of HiC data from juicebox showing partial recue of TADs when CLC and LCL were expressed in the absence of endogenous CTCF. The corresponding FLAG ChIPs are shown on the x-and y-axis. **e** Co-IP experiments showing interaction of RAD21 with transgenic CLC or LCL in the presence (D) and absence (ID) of CTCF.

We next analyzed insulation score at the boundaries of CTCF + CTCFL overlapping sites using the HiCratio method [59]. As expected, boundaries were maximally affected when CTCF was degraded and could be recovered by expression of transgenic CTCF but not CTCFL. In contrast, the presence of transgenic CLC and LCL led to partial recovery of insulation (**Fig. 6c**). Examples of specific sites where partial rescue of TAD structure was achieved by expression of CLC and LCL after removal CTCF are shown in **Fig. 6d.** It is clear in all the above analyses that LCL fared slightly better at rescuing TAD structure and insulation than CLC.

Since the interaction between CTCF and cohesin is important for the establishment of TAD structures [7, 60, 61] we used the chimeric proteins LCL and CLC to determine whether the zinc fingers and/or the N/ C terminal regions were involved. We performed co-immunoprecipitation experiments with lysates from cells harboring chimeric proteins (CLC and LCL) in the presence and absence of CTCF. With this approach we could detect that RAD21 is pulled down with CLC but not LCL in the absence of endogenous CTCF **(Fig. 6e**). This finding demonstrates that the N/C terminal regions of CTCF are involved in mediating the interaction with cohesin. Interestingly, pulldown of RAD21 was observed with LCL in the presence, but not absence of endogenous CTCF. This could be a result of LCL interacting with CTCF as shown by the GFP-Trap experiment (**Additional File 1: Figure S5E**). The failure of LCL to interact with RAD21 in the absence of CTCF indicates that it is the N and/or C terminals of CTCF that mediate interaction with cohesin.

### The N terminus of CTCF interacts with RAD21

To determine whether the N or the C terminal is responsible for CTCF’s ability to interact with cohesin, we inserted transgenic CTCFL chimeric proteins into the *Tigre* locus with either their N or C terminal domains swapped with those of CTCF. The chimeric proteins (**C**TCF N terminus - CTCF**L** zinc fingers - CTCF**L** C-terminus; CTCF**L** N terminus - CTCF**L** zinc fingers - **C**TCF C-terminus) were abbreviated as CLL and LLC where C stands for CTCF and L stands for CTCFL (**Fig. 7a**). The chimeric protein transgenes, along with their FLAG and mRUBY tags, were expressed at the same levels as intact transgenic CTCF and CTCFL, as demonstrated by flow cytometry (**Fig. 7b**). Western blotting confirmed that the level of expression of CLL and LLC were comparable across D and ID conditions (**Fig. 7c**).

**Fig. 7.**
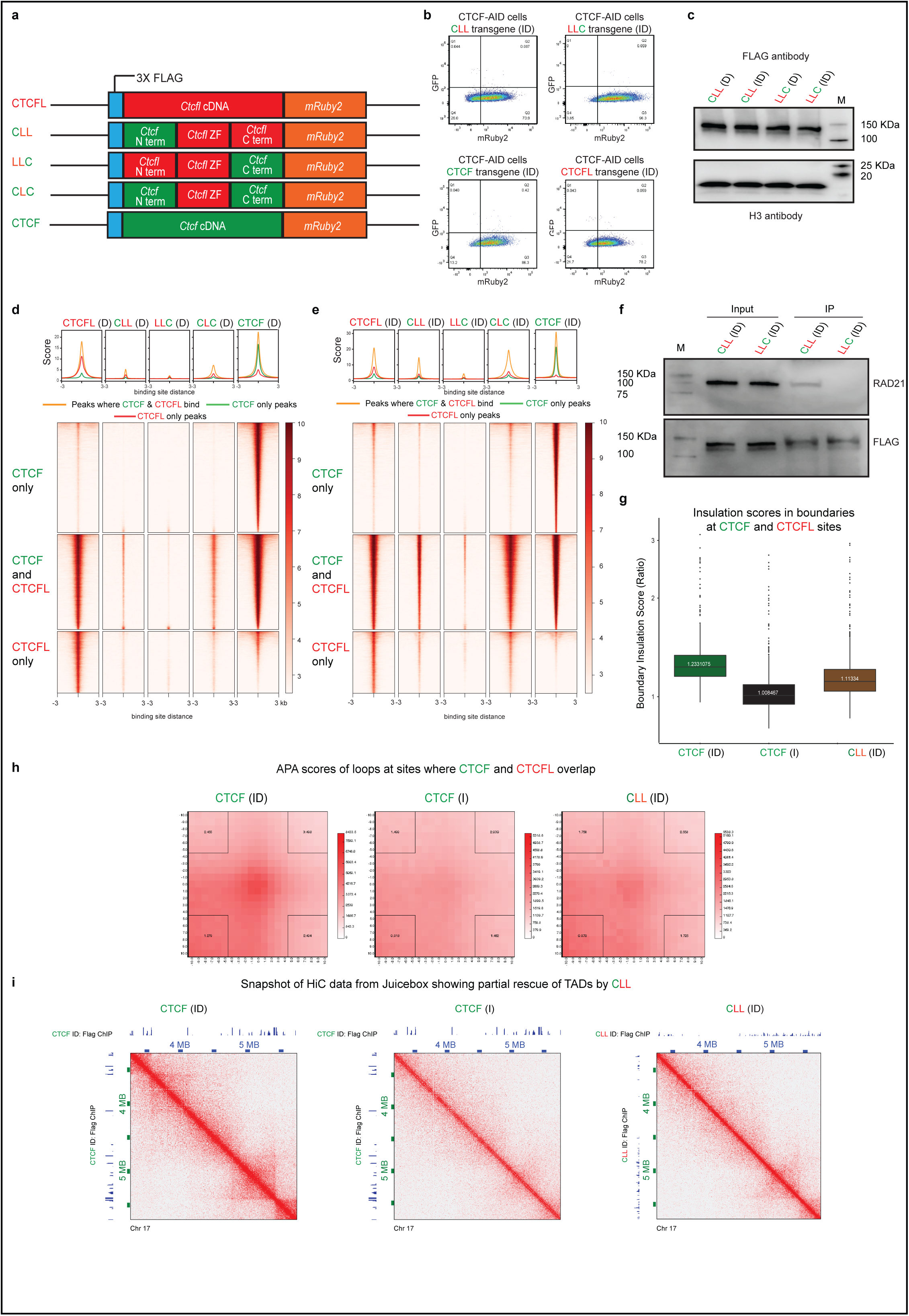
The N terminus of CTCF interacts with RAD21. **a** Schematic showing the doxycycline inducible parent and chimeric protein transgenes knocked into the *Tigre* locus. **b** Flow cytometry confirms that the level of mRuby2 expression of the transgenes and parental proteins are comparable. **c** Western blot using FLAG antibody shows that the level of expression of CLL and LLC is comparable across experimental conditions (D versus ID). Histone H3 serves as loading control. **d,e** Heatmaps showing CTCFL, CLL, LLC, CLC and CTCF ChIP-seq signals at regions where they bind in the presence (**d**) and absence (**e**) of endogenous CTCF. The heatmaps are divided into CTCF only, CTCF + CTCFL overlapping and CTCFL only sites. Average profile of the respective heatmaps are shown above the corresponding heatmaps. **f** Co-IP experiments showing interaction of RAD21 with transgenic CLL and LLC in the absence (ID) of CTCF. M stands for molecular weight marker and the corresponding weights are shown. **g** Insulation scores in boundaries of CTCF + CTCFL overlapping sites in CTCF depleted cells (CTCF-I) and cells depleted of CTCF that were induced to express transgenic CTCF and CLL (ID condition). **h** Aggregate Peak Analysis demonstrates the strength of the loops in the CTCF ID, CTCF I and CLL ID conditions at sites where CTCF and CTCFL bind competitively. **i** Snapshot of HiC data from juicebox showing TADs in presence of transgenic CTCF (CTCF-ID), loss of TADs on CTCF depletion (CTCF-I) and partial recue of TADs when CLL was expressed in the absence of endogenous CTCF (CLL-ID). The corresponding FLAG ChIPs are shown on the x-and y-axis.

FLAG ChIP-seq in the presence of endogenous CTCF (D condition) revealed that CLL has a similar binding profile to CTCFL, although compared to the latter, binding was much reduced. In the case of LLC we detected almost no binding in the Dox condition (**Fig. 7d**). However, binding of both transgenes was increased in the absence of CTCF (ID condition) (**Fig. 7e**), which suggests that the two chimeric proteins are unable to compete effectively with CTCF at these sites. Moreover, these data indicate that the C terminal region of CTCFL is more important than the N terminal region for CTCFL’s binding. Binding of the single swapped chimeric proteins, CLL and LLC was also reduced compared to the double swapped CLC chimera further suggesting that the C and N terminals of CTCF cooperate with each other and are both important for binding.

Co-immunoprecipitation experiments with lysates from cells harboring chimeric protein (CLL) in the presence and absence of CTCF revealed that RAD21 was pulled down with CLL but not with LCC **(Fig. 7f**), indicating that it is the N terminal region of CTCF that is involved in mediating the interaction with cohesin. These interactions are consistent with convergence of CTCF binding being important for loop formation, because the N terminal region of CTCF would be the first point of encounter with cohesin. HiC was performed in cells expressing transgenic CLL in the absence of CTCF to determine if CLL can rescue the chromatin organization lost by CTCF depletion. As seen in **Fig. 7g-i**, CLL was respectively partially able to rescue boundary insulation, aggregate peak enrichment and TAD structure. These results highlight the role played by the N terminus of CTCF in mediating interaction with cohesin as well as genome organization.

## Discussion

CTCF plays a key role in organizing chromatin into highly self-interacting topologically associated domain (TAD) structures by promoting the formation of insulating loops and boundaries that are important for gene regulation. It is a ubiquitously expressed factor in contrast to its paralogue, CTCFL which is normally only transiently present in testis. CTCFL however, is frequently aberrantly expressed in numerous cancers due to genetic abnormalities. As a result of shared and unique zinc finger sequences in CTCF and CTCFL, CTCFL can bind competitively to a subset of CTCF binding sites as well as its own unique locations. While this has been known for some time, the impact of CTCFL on chromosome organization and gene expression has not been comprehensively analyzed in the context of CTCF function. Indeed, CTCFL has largely been studied in a cancer setting which has many other confounding genetic aberrations. Here we made use of a complementation system incorporating auxin inducible degradable endogenous CTCF, combined with doxycycline inducible transgenes encoding CTCF, CTCFL or CTCF-CTCFL chimeric proteins. This approach enabled us to analyze the impact of CTCF and CTCFL expression either individually or in concert, and determine the unique functional impact of each factor as well as the interplay between the two. Use of the chimeric CTCF-CTCFL proteins further provided us with a tool to tease out the contribution of the zinc finger and N/C terminal domains to their individual and shared functions.

Our studies demonstrate that CTCF and CTCFL bind to common and overlapping sites that have distinct properties, highlighting an interesting facet of functional importance: not all CTCF and CTCFL binding sites are created equal. First, CTCF-only binding sites exhibit a preference for intronic and intergenic regions while CTCFL binding is biased towards promoter regions. Specifically, CTCFL is more likely to bind promoter sites than CTCF and when CTCF binds these sites, it prefers locations where CTCF and CTCFL binding overlaps. While there is little overlap in the changes in gene expression mediated by CTCF and CTCFL, of the 219 genes found in the overlapping subset, 146 are regulated by CTCF and/or CTCFL binding to the same promoters and 76 of these are overlapping binding sites. Interestingly, although only 21.52% of CTCF binding events occur at promoter sites, 36.2% of CTCF-mediated gene expression changes are linked to binding at these locations. This implies that, at sites where CTCF binding overlaps with that of CTCFL at promoters, CTCF and CTCFL may act as transcription factors. Alternatively, CTCF might function to bring enhancers closer to promoters by forming loops with CTCF sites present at enhancers. At other sites, namely the predominantly intergenic or intronic CTCF only sites, CTCF may bind enhancers or behave more like an insulator, controlling gene expression in a more distal or indirect manner, respectively.

What mediates the distinct functional impact at the different binding site subsets? We speculate that it has to do with cofactors binding to the C/N terminal domains. Indeed, use of the chimeric proteins allowed us to demonstrate that it is these domains in CTCF and CTCFL that influence promoter versus intronic and intergenic site bias such that the LCL fusion protein has more of a preference for promoter binding sites compared to CTCF and conversely, CLC has more of a preference for intronic and intergenic regions compared to CTCFL. Given that the N and C terminal regions of CTCF and CTCFL are very different it is highly likely that the cofactors they bind are also different. This also likely applies to differences in the zinc fingers between the two proteins. indeed, depletion of RNA has no impact on CTCFL’s binding profile although it has a profound impact on the binding of CTCF as previously shown [38].

As mentioned above, our studies also highlight that CTCFL fails to rescue the insulation boundaries lost on CTCF depletion. We hypothesized that this results from differences in CTCF’s and CTCFL’s relationship with cohesin. In support of this notion, we show that RAD21, a component of the cohesin complex is localized at those sites where CTCF is bound. Furthermore, in contrast to CTCF, CTCFL does not physically interact with cohesin. As a result, CTCFL cannot rescue the effect of CTCF depletion on chromatin folding.

Expression of transgenic CTCF-CTCFL chimeric proteins in the presence and absence of CTCF enabled us to demonstrate that the N terminal region is responsible for CTCF’s interaction with cohesin. Replacing the N terminal region of CTCFL with that of CTCF resulted in partial recovery of TADs, loops and insulation. However, it is of note that LCL did a better job of rescuing all these aspects of chromatin folding after CTCF depletion, despite its inability to interact with cohesin. This underliines the importance of both zinc fingers and N/C terminals in CTCF function.

The RNA Binding regions (RBRs) of ZF1, ZF10 and C terminal regions have been shown to be crucial in binding of CTCF to chromatin, and 3D genome organization [36, 38, 54, 55]. As with the deletion of the RBRs in ZF1 and 10, deletion of the RBR at the C terminal region results in reduced CTCF binding and loss of a subset of CTCF-mediated loops as well as alteration in gene expression [55]. Together these studies indicate that both the C terminal region and zinc fingers contribute to CTCF binding, CTCF-mediated loop formation and gene expression. These findings are consistent with our results showing that the zinc finger and C/N terminal domains have distinct contributions to binding site preference and regulation of chromosome organization. Also consistent with these published studies, we found that the zinc finger and C/N terminal domains have an important functional impact on the transcriptional changes mediated by CTCF and CTCFL. This is highlighted by our finding that overall both CLC and LCL were ineffectual at mediating changes in gene expression in comparison to intact CTCF and CTCFL. However, close inspection of individual genes revealed sites at which the chimeric proteins acted concordantly, discordantly or independent of the parental proteins. Together these findings underscore the combined roles played by the zinc fingers and N/C terminal regions in site-specific regulation of gene expression.

While our manuscript was under revision a paper was published by Pugacheva et al., showing that the N terminal region of CTCF is essential but not sufficient for cohesin retention at CTCF sites [62]. Here the authors examined the binding profiles of cohesin and CTCF but did not demonstrate physical interaction between the two proteins, as we have done here. Their study also demonstrated that CTCFL failed to retain cohesin on chromatin [62]. Although the paper draws some of the same conclusions as ours, the experimental strategy differs in two major ways. First, the cell lines used by Pugacheva et al. expressed an endogenous version of CTCF in which only ZFs 1-8 were functional (ZFs 9-11 mutated), while in our case the auxin inducible degron system allowed us to analyze CTCF and CTCFL chimeric proteins in the presence and absence of endogenous CTCF so we could tease apart their individual effects. Second, their analysis of CTCFL and CTCF-CTCFL chimeric proteins was restricted to the 5,000 sites that either lost or reduced CTCF occupancy when mutant CTCF was expressed, while our study involved a global analysis of CTCF, CTCFL and chimeric proteins.

What role does CTCFL play in regulating chromatin organization and gene expression in a cancer setting where it is expressed in the presence of CTCF? At CTCF + CTCFL overlapping binding sites where CTCFL can bind competitively with CTCF, we demonstrated that even in the presence of CTCF, CTCFL can have an impact on chromosome organization reducing the strength of the aggregate peak enrichment of chromatin loops and in some places abrogating loop formation altogether. Importantly, binding of CTCFL at CTCF + CTCFL overlapping binding sites was linked to differential expression of genes within the loops. These findings have implications for the role of CTCFL in altering chromatin organization and gene expression in the context of cancer.

## Conclusion

In sum, use of the complementation system incorporating auxin degradable endogenous CTCF combined with doxycycline inducible transgenic CTCF, CTCFL and CTCF-CTCFL in the presence and absence of CTCF enabled us to demonstrate that CTCF’s and CTCFL’s unique and overlapping binding sites have distinct binding sequences, biases for being in promoters rather than intronic or intergenic regions and effects on chromatin folding. Furthermore, our studies highlight unique functional aspects of the zinc finger and C/N terminal domains of CTCF and CTCFL in controlling binding site preference as well as site-specific effects on chromosome organization and gene expression. Future studies will clarify the identity of the cofactors that facilitate the site-specific functions of CTCF and CTCFL, and the genetic system we have developed here will be a useful tool for addressing this question.

## Methods

### Cell lines

Mouse embryonic stem cells E14Tg2a (karyotype 19, XY; 129/Ola isogenic background) and all clones derived from these were cultured under feeder-free conditions in 0.1% gelatin (Sigma ES- 006-B) coated dishes (Falcon, 353003) at 37°C and 5% CO_2_ in a humidified incubator. The cells were grown in DMEM (Thermo Fisher, 11965-118) supplemented with 15% fetal bovine serum (Thermo Fisher, SH30071.03), 100□U/ml penicillin - 100□μg/ml streptomycin (Sigma, P4458), 1 X GlutaMax supplement (Thermo Fisher, 35050-061), 1 mM sodium pyruvate (Thermo Fisher, 11360- 070), 1 X MEM non-essential amino-acids (Thermo Fisher, 11140-50), 50 μM b-mercaptoethanol (Sigma, 38171), 10^4^ U/ml leukemia inhibitory factor (Millipore, ESG1107), 3□μM CHIR99021 (Sigma, SML1046) and 1□μM MEK inhibitor PD0325901 (Sigma, PZ0162). The cells were passaged every alternate day by dissociation with TrypLE (Thermo Fisher, 12563011).

### DNA constructs

#### Construction of vector for cloning transgenic, doxycycline-inducible expression of *Ctcfl*

cDNA clone for *Mus musculus Ctcfl* (NCBI Gene ID: 664799) was purchased from Transomic Technologies (TCMS1004). The cDNA was amplified such that it harbors AflII sequence at the 3’ end of the gene and was fused with FLAG tag (that harbors NotI sequence) at 5’ end with the help of a fusion PCR. The resultant fragment was digested with NotI and AflII. The *Ctcf* gene was removed from pEN366 [7] by digesting with the same enzymes. This backbone was used for insertion of *Ctcfl* as well as the chimeric constructs.

#### For construction of *Ctcf* and *Ctcfl* with the terminals swapped

To construct a hybrid gene with *Ctcf* N terminus - *Ctcfl* zinc fingers - *Ctcf* C-terminus, the region encoding the first 265 amino acids of mice *Ctcf* was fused in frame to the region encoding amino acids 259 to 568 of mice *Ctcfl* and the 159 (578-736) C-terminal amino acids of *Ctcf*. The fragments of *Ctcf* were amplified from pEN366 [7] and *Ctcfl* from cDNA clone (TCMS1004, Transomic Technologies). The resulting plasmid was named pCLC (‘C’ for *Ctcf* and ‘L’ for *Ctcfl*) and the transgene is referred to as CLC henceforth. Similarly, to construct a hybrid of mice *Ctcfl* N terminus - *Ctcf* zinc fingers - *Ctcfl* C-terminus protein, the region encoding the first 258 amino acids of *Ctcfl* was fused in frame to the regions encoding amino acids 266 to 577 of *Ctcf* and the 68 (569-636) C-terminal amino acids of *Ctcfl*. The plasmid was named pLCL and the transgene as LCL respectively. The construction of these mutant genes was achieved by swapping one terminus at a time using a two-step PCR overlap extension method. In brief, cDNA region corresponding to each of the terminals and zinc fingers were PCR amplified in such a way that it included a short stretch of the 5′ and/or 3′ region of the neighboring fragment to be connected. The desired PCR products were then annealed, amplified by PCR and cloned into the NotI and AflII sites of pEN366 backbone. All of the constructs were verified by DNA sequence analysis. The transgenes with one terminus each of CTCFL swapped with that of CTCF were constructed and named using the same terminology as LLC (*Ctcfl* with C-terminal *Ctcf*) and CLL (*Ctcfl* with N-terminal *Ctcf*). With all transgenes, the final vector harbors an N terminal 3 X FLAG tag and a C terminal *mRuby* as in-frame fusion to the transgenes (*Ctcfl, Ctcf*, LCL, CLC, LLC and CLL). It also harbors *TetO-3G* element and *rtTA3G* for doxycycline induced expression of the transgene and homology arms surrounding the sgRNA target site of the *Tigre* locus for locus-specific insertion. The selection of stable integrants was achieved by virtue of *FRT-PGK-puro-FRT* cassette. Further details of the vector are described elsewhere [7]. The vector pX330-EN1201 [7] harboring spCas9 nuclease and sgRNAs was used for targeting of the *Tigre* locus.

A table listing the transgenes, treatment conditions, experiments performed and citing figures is provided in **Additional File 4: Table S3**.

### Gene targeting

Mouse embryonic stem cell E14Tg2a harboring *Ctcf-AID-eGFP* on both alleles and a knock-in of pEN114 - *pCAGGS-Tir1-V5-BpA-Frt-PGK-EM7-PuroR-bpA-Frt-Rosa26* at *Rosa26* locus was used as the parental cell line for making all the transgenes [7]. pEN366 derived vectors harboring the rescue transgenes (*Ctcf, Ctcfl* as well as chimeric proteins) were used for targeting transgenes to the *Tigre* locus [7]. For nucleofections, 15 μg each of plasmids harboring the transgenes and 2.5 μg of those with sgRNA targeting the *Tigre* locus was used. Nucleofection were performed using Amaxa P3 Primary Cell kit (Lonza, V4XP-3024) and 4D- transfector. 2 million cells were transfected with program CG-104 in each case. The cells were recovered for 48 h with no antibiotic followed by selection in puromycin (1 μg/mL) (Thermo Fisher, A1113803). Single colonies were manually picked and expanded in 96 well plates. Clones were genotyped by PCR and FACS was performed to confirm that the level of expression of transgenes were comparable. All the clones that were used for the analyses were homozygous for the integration of the transgenes and their levels of expression were comparable.

### Induction of auxin inducible degradation of CTCF and doxycycline induced expression

For degradation of endogenous CTCF, the auxin-inducible degron was induced by adding 500 μM indole-3-acetic acid (IAA, chemical analog of auxin) (Sigma, I5148) to the media. Expression of transgenes were achieved by the addition of doxycycline (Dox, 1 μg/ml) (Sigma, D9891) to the media. The cells were treated with IAA and/or Dox for 2 days unless mentioned otherwise.

### Western Blotting

mESCs were dissociated using TrypLE, washed in PBS, pelleted and used for western blotting. Approximately 2 million cells were used to prepare cell extract. Cell pellets were resuspended in RIPA lysis buffer (Thermo Fisher, 89900) with 1X HALT protease inhibitors (Thermo Fisher, 78430), incubated on ice for 30 min, spun at 4°C at 13,000 rpm for 10 min and supernatant was collected. For the western blot of CTCF, low salt lysis buffer (0.1 M NaCl, 25 mM HEPES, 1 mM MgCl_2_, 0.2 mM EDTA and 0.5% NP40) was used supplemented with 125 U/ml of benzonase (Sigma E1014). Protein concentration was measured using the Pierce BCA assay kit (Thermo Fisher, 23225). 20 μg of protein were mixed with Laemmli buffer (Biorad, 1610737) and b-mercaptoethanol, heated at 95°C for 10 min and run on a Mini-protean TGX 4%-20% polyacrylamide gel (Biorad, 456-1095). The proteins were transferred onto PVDF membranes using the Mini Trans-Blot Electrophoretic Transfer Cell (Bio-Rad, 170-3930) at 80 V, 120 mA for 90 min. PVDF membranes were blocked with 5% BSA in 1 X TBST prior to the addition of antibody. The membranes were probed with appropriate antibodies overnight at 4°C (anti-rabbit histone H3 (abcam, ab1791; 1: 2,500 dilution), anti-mouse FLAG antibody (Sigma, F1804; 1: 1,000 dilution), anti CTCF (Active Motif, 61311), anti Rad21 (ab992) and anti-GFP (Abcam, ab6556)). Membranes were washed five times in PBST (1 × PBS and 0.1% Tween 20) for 5 min each and incubated with respective secondary antibodies in 5% BSA at room temperature for 1□h. The blots were rinsed in PBST and developed using enhanced chemiluminescence (ECL) and imaged by Odyssey LiCor Imager (Kindle Bioscien ces).

### RNase A treatment

Briefly, mESCs were collected following trypsinsation, washed twice in PBS, permeabilized with 0.05% Tween-20 in PBS for 10 min on ice, washed, resuspended in PBS and incubated with RNase A (1 mg/ml) for 30 minutes at room temperature [63]. Cells were washed twice in PBS and cross linked for ChIPmentation with FLAG antibody.

### Immunoprecipitation

For immunoprecipitation of nuclear lysates, cells were first lysed in 5 X pellet-volume of ice-cold Buffer A (10 mM Tris-HCl (pH 7.5-7.9), 1.5 mM MgCl_2_, 10 mM KCl, 0.5 mM DTT, 0.2 mM PMSF, 0.1% NP40) supplemented with complete EDTA-free tablets (Roche) while rotating in the cold-room for 10 minutes. Nuclei fractions were then isolated by spinning down the lysate at 1000 xg for 5 minutes at 4°C. The remaining nuclear pellet was then resuspendded in 5 X pellet-volume of ice-cold Buffer C (10 mM Tris-HCl (pH 7.5-7.9), 25% glycerol, 0.42 M NaCl, 1.5 mM MgCl_2_, 0.2 mM EDTA, 0.5 mM DTT, 0.5 mM PMSF) supplemented with complete EDTA-free tablets and placed on the cold-room rotator for 120 minutes. Soluble nuclear extracts were then cleared by centrifugation at 20,000 xg for 10 minutes at 4°C. The remaining insoluble nuclear pellet was dissolved in 3 X pellet-volume of Urea-Chaps Buffer (8 M Urea, 20 mM HEPES, 1% CHAPs), supplemented with 1 X Halt Protease (ThermoFisher, 87786), vortexed vigorously at 10-minute intervals over a 30- minute incubation at room-temperature, and then combined with the soluble nuclear extract to make the complete nuclear lysate. BCA Assay (ThermoFisher, 23225) was used to determine protein levels of each sample in which 2 mg of nuclear lysates was incubated overnight with 50 uL of ANTI-FLAG M2 magnetic beads (Sigma Cat# M8823) at 4°C or for 2 hours with 20 uL of GFP-Trap magnetic beads (Chromotek, gtma-20) at 4°C. Beads were washed 3 X in ice-cold IP wash buffer (20 mM Tris-HCl (pH 7.5-7.9), 150 mM NaCl, 1 mM EDTA, 0.05% Triton X-100, 5% Glycerol). FLAG immunoprecipitates were eluted at 95°C for 10 minutes and GFP immunoprecipitates were eluted at 60°C for 20 minutes into 1 X SDS-Page Buffer (Bio-Rad) supplemented with 5% BME. Samples were resolved by SDS-Page using 4-20% gradient gels (BioRad) and transferred to PVDF membranes by a wet transfer protocol. Immunoblotting was performed using 5% BSA for both blocking and primary or secondary horseradish peroxidase-conjugated antibody incubation. Primary antibodies used were anti-FLAG M2 (Sigma, F1804) (1:1000), anti-Rad21 (Abcam, ab992) (1:1000) or anti-GFP (Abcam, ab6556) (1:1000) and secondary antibodies used were Mouse IgG HRP Linked Whole (Sigma, GENA931) (1:2000), Mouse Anti-Rabbit IgG (Light Chain Specific) (CST, #93702S) (1:5000) or Rabbit IgG HRP Linked Whole (GE Healthcare, NA9340), respectively. Blots were developed using enhanced chemiluminescence (ECL) and imaged by Odyssey LiCor Imager (Kindle Biosciences).

### Flow cytometric analysis

Cells were dissociated with TrypLE, washed and resuspended in MACS buffer for flow cytometric analysis on LSRII UV (BD Biosciences). Analysis was performed using the FlowJo software.

### Microscopy

Images were acquired on EVOS FL Color Imaging System using a 20 X objective.

### ChIPmentation

mESCs were dissociated using TrypLE, washed in PBS and fixed in 1% formaldehyde for 10 min at room temperature. Quenching was performed by adding glycine to a final concentration of 0.125 M followed by incubations of 5 min at room temperature and 15 min at 4°C. The cells were washed twice in PBS with 0.125 M glycine, pelleted, snap frozen and stored at −80°C till use. Fixed cells (10 million) were thawed on ice, resuspended in 350 μl ice cold lysis buffer (10 mM Tris-HCl (pH 8.0), 100 mM NaCl, 1 mM EDTA (pH 8.0), 0.5 mM EGTA (pH 8.0), 0.1% sodium deoxycholate, 0.5% N- lauroysarcosine and protease inhibitors) and lysed for 10 min by rotating at 4°C. Chromatin was sheared using a bioruptor (Diagenode) (25 cycles: 30 sec on, 30 sec off). Triton X-100 was added to a final concentration of 1% and the samples were centrifuged for 5 min at 16000 rcf at 4°C. Supernatant was collected and shearing was continued for another 10 min and the chromatin was quantified. FLAG M2 Magnetic Beads (Sigma, M8823) were used for FLAG ChIPs. In other cases (CTCF, Cohesin, IgG) antibodies were bound to protein A magnetic beads by incubation on a rotator for one hour at room temperature. 10 μl each of antibody was bound to 50 μl of protein-A magnetic beads (Dynabeads). and added to the sonicated chromatin from 10 million cells per immunoprecipitation. The beads were washed and tagmentation were performed as per the original ChIPmentation protocol (Schmidl et al., 2015). In short, the beads were washed twice in 500 μl cold low-salt wash buffer (20 mM Tris-HCl (pH 7.5), 150 mM NaCl, 2 mM EDTA (pH 8.0), 0.1% SDS, 1% tritonX-100), twice in 500 μl cold LiCl-containing wash buffer (10 mM Tris-HCl (pH 8.0), 250 mM LiCl, 1 mM EDTA (pH 8.0), 1% triton X-100, 0.7% sodium deoxycholate) and twice in 500 μl cold 10 mM cold Tris-Cl (pH 8.0) to remove detergent, salts and EDTA. Subsequently, the beads were resuspended in 25 μl of the freshly prepared tagmentation reaction buffer (10 mM Tris-HCl (pH 8.0), 5 mM MgCl2, 10% dimethylformamide) and 1 μl Tagment DNA Enzyme from the Nextera DNA Sample Prep Kit (Illumina) and incubated at 37°C for 1 min in a thermocycler. Following tagmentation, the beads were washed twice in 500 μl cold low-salt wash buffer (20 mM Tris-HCl (pH 7.5), 150 mM NaCl, 2 mM EDTA (pH8.0), 0.1% SDS, 1% triton X-100) and twice in 500 μl cold Tris-EDTA-Tween buffer (0.2% tween, 10 mM Tris-HCl (pH 8.0), 1 mM EDTA (pH 8.0)). Chromatin was eluted and de-crosslinked by adding 70 μl of freshly prepared elution buffer (0.5% SDS, 300 mM NaCl, 5 mM EDTA (pH 8.0), 10 mM Tris-HCl (pH 8.0) and 10 ug/ml proteinase K for 2 hours at 55°C and overnight at 65°C. The supernatant was collected and saved. The beads were supplemented with an additional 30 μl of elution buffer, incubated for 1 h at 55°C and the supernatants were combined. DNA was purified using MinElute Reaction Cleanup Kit (Qiagen 28204) and eluted in 20 ul. Purified DNA (20 μl) was amplified as per the ChIPmentation protocol [64] using indexed and non-indexed primers and NEBNext High-Fidelity 2X PCR Master Mix (NEB M0541) in a thermomixer with the following program: 72°C for 5 m; 98°C for 30 s; 14 cycles of 98°C for 10 s, 63°C for 30 s, 72°C for 30 s and a final elongation at 72°C for 1 m. DNA was purified using Agencourt AMPure XP beads (Beckman, A63881) to remove fragments larger than 700 bp as well as the primer dimers. Library quality and quantity were estimated using Tapestation bioanalyzer (Agilent) as well as Qubit (ThermoFisher) assays. Samples were quantified using and Library Quantification Kit (Kapa Biosystems, KK4824) and sequenced with Illumina Hi-Seq 4000 using 50 cycles single-end mode.

### RNA seq

mESCs were dissociated using TrypLE, washed in PBS, pelletted and used used for extracting RNA. RNA from were extracted from 2.5 million cells using RNeasy plus kit (Qiagen 74134) in each case. The poly-adenylated transcripts were positively selected from the RNA using the NEBNext Poly(A) mRNA Magnetic Isolation Module (E7490) following the manufacturer’s protocol. Libraries were prepared according to the directional RNA-seq dUTP method adapted from http://wasp.einstein.yu.edu/index.php/Protocol:directionalWholeTranscript_seq that preserves information about transcriptional direction. Library concentrations were estimated using tapestation and Qubit assays, pooled and sequenced on a Next-seq instrument (Illumina Hi-Seq 4000) using 50 cycles paired-end mode.

### Hi-C

Hi-C was performed in duplicate using 1 million cells each. MESCs were dissociated using TrypLE, washed in PBS and fixed in 1% formaldehyde for 10 min at room temperature. Quenching was performed by adding glycine to a final of 0.125 M followed by incubations of 5 min at room temperature and 15 min at 4°C. Hi-C samples were processed using the Arima Hi-C kit as per the manufacturer’s protocol and sequenced with Illumina NovaSeq 6000 using 50 cycles paired-end mode.

## QUANTIFICATION AND STATISTICAL ANALYSIS

### ChIP-seq data processing and quality control

Reads were aligned to GRCm38/mm10 genome with Bowtie2 [65] (parameters: –no-discordant -p 12 –no-mixed -N 1 -X 2000). Ambiguous reads were filtered to use uniquely mapped reads in the downstream analysis. PCR duplicates were removed using Picard-tools (version 1.88). For FLAG and RAD21 ChIP-seq, MACS version 1.4.2 [66] was used to call peaks (parameters: -g 1.87e9 -- qvalue 0.05 for FLAG; --broad -q 0.05 for RAD21). Bigwigs were obtained for visualization on individual as well as merged bam files using Deeptools/2.3.3 [67] (parameters: bamCoverage -- binSize 1 --normalizeUsing RPKM). Heatmaps and average profiles were performed on merged bigwig files using Deeptools/2.3.3. We also used DiffBind package [68] to cluster the samples and generate heatmaps (Parameters: summits=250).

### RNA-seq data processing and quality control

Raw sequencing files were aligned against the mouse reference genome (GRCm38/mm10) using the STAR [69] aligner (v.2.6) and differentially expressed genes were called using DESeq2 [70] with an adjusted p-value of 0.01 and a fold change cutoff of 1. Venn diagrams were generated using the ‘eulerr’ [71] library in R package. We obtained a list of mouse TSS coordinates from the Ensembl database (GRCm38.p6 - release 98) [72] that was used in the downstream analyses.

### Hi-C Processing and Quality Control

Hi-C-Bench [73] was used to align and filter the Hi-C data and identify TADs. To generate Hi-C filtered contact matrices, the Hi-C reads were aligned against the mouse reference genome (GRCm38/mm10) by bowtie2 (version 2.3.1). Mapped read pairs were filtered by the GenomicTools tools-hic filter command integrated in HiC-bench for known artifacts of the Hi-C protocol. The filtered reads include multi-mapped reads (‘multihit’), read-pairs with only one mappable read (‘single sided’), duplicated read-pairs (‘ds.duplicate’), low mapping quality reads (MAPQ < 30), read-pairs resulting from self-ligated fragments, and short-range interactions resulting from read-pairs aligning within 25kb (‘ds.filtered’). For the downstream analyses, all the accepted intra-chromosomal read-pairs (‘ds.accepted intra’) were used. The total numbers of reads in the 2 biological replicates for each condition ranged from ∼130 million reads to ∼300 million. The percentage of reads aligned was always over 97% in all samples. The proportion of accepted reads (‘ds-accepted-intra’ and ‘ds-accepted-inter’) was ∼40%, which in all cases was sufficient to annotate TADs with HiC-Bench.

## DOWNSTREAM ANALYSIS

### Annotation of ChIP peak sets and motif analysis

To obtain a peak set per condition we first merged the peaks in each replicate (overlap ≥ 1 bp) and then only the peaks present in both replicates were considered (overlap ≥ 1 bp). ‘CTCF-only’ sites correspond to peaks present in the FLAG ChIP seq of CTCF (ID) peak set and absent in the CTCFL (D) set. The ‘CTCF and CTCFL sites’ has the peaks that were found in both, CTCF (ID) and CTCFL (D) peak sets. ‘CTCFL-only’ sites correspond to peaks present in the CTCFL (D) peak set and absent in the CTCF (ID) set. A peak was considered present in two conditions when the peak overlap was higher than 66% for both peaks. We used the ChIPSeeker [75] library to annotate the peak sets obtained. Annotation packages: ‘TxDb.Mmusculus.UCSC.mm10.knownGene’ and ‘org.Mm.eg.db’ (Bioconductor). Promoters were defined as ± 3 kilobases from the transcription start site. Venn diagrams were generated using the ‘bedr’ library [75] in R package. The MEME-ChIP tool from the MEME suite [76, 77] was used to detect motifs in the peak sets.

### Compartments, TADs and Boundaries

#### Compartment Analysis

Compartment analysis was carried out using the HOMER [78] pipeline (v4.6). Hi-C filtered matrices were given as input together with ATAC-seq peaks for compartment prediction (default parameters: 50 kb resolution). HOMER was used to perform a principal component analysis of the normalized interaction matrices and then, we used the PCA1 component to predict regions of active (A compartments) and inactive chromatin (B compartments), and to generate the eigenvalues bedgraph files of each condition. HOMER assumes that gene-rich regions with active chromatin marks have similar PC1 values, while gene deserts show differing PC1 values.

#### Domain boundary Insulation Scores

The Hi-C filtered contact matrices were corrected using the ICE “correction” algorithm [79] built into HiC-bench. Chromatin domains and boundaries were called using Hicratio [59] at 40 kb resolution. We also called domains using the Crane algorithm [80] at 40 kb bin resolution with an insulating window of 500 kb. Hi-C heatmaps for regions of interest were generated in Juicebox [81]. To assess and compare boundary strength alteration across all the conditions using the HiCratio method, we calculated insulation score for each 40 kb resolution bin, as described by Lazaris et al. [73]. Then, TAD boundaries of size 40 kb were identified as local maxima of the insulation scores. Only insulation scores above a certain cutoff were considered as potential TAD boundaries. We determined the false discovery rate by repeating the same analysis on perturbed matrices. TAD boundaries were reported at 5% false discovery rate. Insulation scores for all conditions were matched to every boundary identified in the CTCFL (U) condition (reference boundaries).

#### Loop Analysis

Loops were annotated for all conditions using HiCCUPS [82]. Loops were called at 25 kb resolution using default parameters (KR-normalization). We also assessed chromatin loops by using the aggregate peak analysis (APA) at 10 kb resolution (-r 10000 -k KR).

#### Availability of data and materials

1. All raw and processed sequencing data files have been deposited at NCBI’s Gene Expression Omnibus (GEO) accession GSE GSE140363 (84).

## Ethics approval and consent to participate

Not applicable

## Supplementary Information

**Additional File 1:**

**Supplementary figures S1 to S7**

**Additional File 2**

**Table S1: Differentially expressed genes after CTCF degradation and CTCFL induction.** Genes upregulated and downregulated under conditions, CTCF U vs I, CTCFL U vs D and CTCFL U vs ID are listed. Log_2_ fold change and adjusted p values are given in each case.

**Additional File 3**

**Table S2: HiC-bench filter stats.** The filter stats of HiC experiments using all transgenes (CTCF, CTCFL, CLC, LCL) under conditions of U, I, D and ID.

**Additional File 4**

**Table S3: Information regarding experiments performed with the transgenes under different conditions and the figures where the relevant data is shown.**

## Author’ contributions

Conceptualization & Study Design, N.M, J.S; Formal analysis, J-R.H, A.R and S.B; Writing – Original Draft, N.M and J.S; Supervision, A.T and J.S

## Acknowledgements

The authors thank Skok lab members for helpful scientific discussions, New York University School of Medicine High Performance Computing Facility (HPCF) for computing technical support, Adriana Heguy and the Genome Technology Center (GTC) core for sequencing efforts, Applied Bioinformatics Laboratories (ABL) for providing bioinformatics support and helping with the analysis and interpretation of the data and the NYU Flow Cytometry and Cell Sorting Center for FACS analysis and sorting. GTC and ABL are shared resources partially supported by the Cancer Center Support Grant P30CA016087 at the Laura and Isaac Perlmutter Cancer Center.

## References

1. Tang Z, Luo OJ, Li X, Zheng M, Zhu JJ, Szalaj P, Trzaskoma P, Magalska A, Wlodarczyk J, Ruszczycki B, et al: CTCF-Mediated Human 3D Genome Architecture Reveals Chromatin Topology for Transcription. Cell 2015, 163: 1611–1627.

2. Sanborn AL, Rao SS, Huang SC, Durand NC, Huntley MH, Jewett AI, Bochkov ID, Chinnappan D, Cutkosky A, Li J, et al: Chromatin extrusion explains key features of loop and domain formation in wild-type and engineered genomes. Proc Natl Acad Sci U S A 2015, 112: E6456–6465.

3. Nuebler J, Fudenberg G, Imakaev M, Abdennur N, Mirny LA: Chromatin organization by an interplay of loop extrusion and compartmental segregation. Proc Natl Acad Sci U S A 2018, 115: E6697–E6706.

4. Nasmyth K: Disseminating the genome: joining, resolving, and separating sister chromatids during mitosis and meiosis. Annu Rev Genet 2001, 35: 673–745.

5. Fudenberg G, Imakaev M, Lu C, Goloborodko A, Abdennur N, Mirny LA: Formation of chromosomal domains by loop extrusion. Cell reports 2016, 15: 2038–2049.

6. Gomez-Marin C, Tena JJ, Acemel RD, Lopez-Mayorga M, Naranjo S, de la Calle-Mustienes E, Maeso I, Beccari L, Aneas I, Vielmas E, et al: Evolutionary comparison reveals that diverging CTCF sites are signatures of ancestral topological associating domains borders. Proc Natl Acad Sci U S A 2015, 112: 7542–7547.

7. Nora EP, Goloborodko A, Valton A-L, Gibcus JH, Uebersohn A, Abdennur N, Dekker J, Mirny LA, Bruneau BG: Targeted degradation of CTCF decouples local insulation of chromosome domains from genomic compartmentalization. Cell 2017, 169: 930-944. e922.

8. Loukinov DI, Pugacheva E, Vatolin S, Pack SD, Moon H, Chernukhin I, Mannan P, Larsson E, Kanduri C, Vostrov AA: BORIS, a novel male germ-line-specific protein associated with epigenetic reprogramming events, shares the same 11-zinc-finger domain with CTCF, the insulator protein involved in reading imprinting marks in the soma. Proceedings of the National Academy of Sciences 2002, 99: 6806–6811.

9. Hore TA, Deakin JE, Graves JAM: The evolution of epigenetic regulators CTCF and BORIS/CTCFL in amniotes. PLoS genetics 2008, 4: e1000169.

10. Sleutels F, Soochit W, Bartkuhn M, Heath H, Dienstbach S, Bergmaier P, Franke V, Rosa-Garrido M, van de Nobelen S, Caesar L: The male germ cell gene regulator CTCFL is functionally different from CTCF and binds CTCF-like consensus sites in a nucleosome composition-dependent manner. Epigenetics & chromatin 2012, 5: 8.

11. Soltanian S, Dehghani H: BORIS: a key regulator of cancer stemness. Cancer cell international 2018, 18: 1–13.

12. Suzuki T, Kosaka-Suzuki N, Pack S, Shin D-M, Yoon J, Abdullaev Z, Pugacheva E, Morse HC, Loukinov D, Lobanenkov V: Expression of a testis-specific form of Gal3st1 (CST), a gene essential for spermatogenesis, is regulated by the CTCF paralogous gene BORIS. Molecular and cellular biology 2010, 30: 2473–2484.

13. Bhan S, Negi SS, Shao C, Glazer CA, Chuang A, Gaykalova DA, Sun W, Sidransky D, Ha PK, Califano JA: BORIS binding to the promoters of cancer testis antigens, MAGEA2, MAGEA3, and MAGEA4, is associated with their transcriptional activation in lung cancer. Clinical cancer research 2011, 17: 4267–4276.

14. Hong JA, Kang Y, Abdullaev Z, Flanagan PT, Pack SD, Fischette MR, Adnani MT, Loukinov DI, Vatolin S, Risinger JI: Reciprocal binding of CTCF and BORIS to the NY-ESO-1 promoter coincides with derepression of this cancer-testis gene in lung cancer cells. Cancer research 2005, 65: 7763–7774.

15. Kang Y, Hong J, Chen G, Nguyen D, Schrump D: Dynamic transcriptional regulatory complexes including BORIS, CTCF and Sp1 modulate NY-ESO-1 expression in lung cancer cells. Oncogene 2007, 26: 4394.

16. D’Arcy V, Abdullaev ZK, Pore N, Docquier F, Torrano V, Chernukhin I, Smart M, Farrar D, Metodiev M, Fernandez N: The potential of BORIS detected in the leukocytes of breast cancer patients as an early marker of tumorigenesis. Clinical cancer research 2006, 12: 5978–5986.

17. D’arcy V, Pore N, Docquier F, Abdullaev Z, Chernukhin I, Kita G, Rai S, Smart M, Farrar D, Pack S: BORIS, a paralogue of the transcription factor, CTCF, is aberrantly expressed in breast tumours. British journal of cancer 2008, 98: 571.

18. Risinger JI, Chandramouli GV, Maxwell GL, Custer M, Pack S, Loukinov D, Aprelikova O, Litzi T, Schrump DS, Murphy SK: Global expression analysis of cancer/testis genes in uterine cancers reveals a high incidence of BORIS expression. Clinical cancer research 2007, 13: 1713–1719.

19. Okabayashi K, Fujita T, Miyazaki J, Okada T, Iwata T, Hirao N, Noji S, Tsukamoto N, Goshima N, Hasegawa H: Cancer-testis antigen BORIS is a novel prognostic marker for patients with esophageal cancer. Cancer science 2012, 103: 1617–1624.

20. He J, Huang Y, Liu Z, Zhao R, Liu Q, Wei L, Yu X, Li B, Qin Y: Hypomethylation of BORIS is a promising prognostic biomarker in hepatocellular carcinoma. Gene 2017, 629: 29–34.

21. Hillman JC, Pugacheva EM, Barger CJ, Sribenja S, Rosario S, Albahrani M, Truskinovsky AM, Stablewski A, Liu S, Loukinov DI: BORIS expression in ovarian cancer precursor cells alters the CTCF cistrome and enhances invasiveness through GALNT14. Molecular Cancer Research 2019, 17: 2051–2062.

22. Salgado-Albarrán M, González-Barrios R, Guerra-Calderas L, Alcaraz N, Sánchez-Correa TE, Castro-Hernández C, Sánchez-Pérez Y, Aréchaga-Ocampo E, García-Carrancá A, de León DC: The epigenetic factor BORIS (CTCFL) controls the androgen receptor regulatory network in ovarian cancer. Oncogenesis 2019, 8: 1–12.

23. Woloszynska-Read A, James SR, Link PA, Yu J, Odunsi K, Karpf AR: DNA methylation-dependent regulation of BORIS/CTCFL expression in ovarian cancer. Cancer Immunity Archive 2007, 7: 21.

24. Woloszynska-Read A, Zhang W, Yu J, Link PA, Mhawech-Fauceglia P, Collamat G, Akers SN, Ostler KR, Godley LA, Odunsi K: Coordinated cancer germline antigen promoter and global DNA hypomethylation in ovarian cancer: association with the BORIS/CTCF expression ratio and advanced stage. Clinical cancer research 2011, 17: 2170–2180.

25. Cheema Z, Hari-Gupta Y, Kita GX, Farrar D, Seddon I, Corr J, Klenova E: Expression of the cancer-testis antigen BORIS correlates with prostate cancer. The Prostate 2014, 74: 164–176.

26. Hoffmann MJ, Müller M, Engers R, Schulz WA: Epigenetic control of CTCFL/BORIS and OCT4 expression in urogenital malignancies. Biochemical pharmacology 2006, 72: 1577–1588.

27. Debruyne DN, Dries R, Sengupta S, Seruggia D, Gao Y, Sharma B, Huang H, Moreau L, McLane M, Day DS: BORIS promotes chromatin regulatory interactions in treatment-resistant cancer cells. Nature 2019, 572: 676–680.

28. Klenova EM, Morse III HC, Ohlsson R, Lobanenkov VV: The novel BORIS+ CTCF gene family is uniquely involved in the epigenetics of normal biology and cancer. In Seminars in cancer biology. Elsevier; 2002: 399–414.

29. Pugacheva EM, Suzuki T, Pack SD, Kosaka-Suzuki N, Yoon J, Vostrov AA, Barsov E, Strunnikov AV, Morse III HC, Loukinov D: The structural complexity of the human BORIS gene in gametogenesis and cancer. PloS one 2010, 5: e13872.

30. Cheever MA, Allison JP, Ferris AS, Finn OJ, Hastings BM, Hecht TT, Mellman I, Prindiville SA, Viner JL, Weiner LM: The prioritization of cancer antigens: a national cancer institute pilot project for the acceleration of translational research. Clinical cancer research 2009, 15: 5323–5337.

31. Hashimoto H, Wang D, Horton JR, Zhang X, Corces VG, Cheng X: Structural Basis for the Versatile and Methylation-Dependent Binding of CTCF to DNA. Mol Cell 2017.

32. Yin M, Wang J, Wang M, Li X, Zhang M, Wu Q, Wang Y: Molecular mechanism of directional CTCF recognition of a diverse range of genomic sites. Cell research 2017, 27: 1365.

33. Nakahashi H, Kwon K-RK, Resch W, Vian L, Dose M, Stavreva D, Hakim O, Pruett N, Nelson S, Yamane A: A genome-wide map of CTCF multivalency redefines the CTCF code. Cell reports 2013, 3: 1678–1689.

34. Marshall AD, Bailey CG, Rasko JE: CTCF and BORIS in genome regulation and cancer. Current opinion in genetics & development 2014, 24: 8–15.

35. Pugacheva EM, Rivero-Hinojosa S, Espinoza CA, Méndez-Catalá CF, Kang S, Suzuki T, Kosaka-Suzuki N, Robinson S, Nagarajan V, Ye Z: Comparative analyses of CTCF and BORIS occupancies uncover two distinct classes of CTCF binding genomic regions. Genome biology 2015, 16: 161.

36. Saldaña-Meyer R, González-Buendía E, Guerrero G, Narendra V, Bonasio R, Recillas-Targa F, Reinberg D: CTCF regulates the human p53 gene through direct interaction with its natural antisense transcript, Wrap53. Genes & development 2014, 28: 723–734.

37. Xiao T, Wallace J, Felsenfeld G: Specific sites in the C terminus of CTCF interact with the SA2 subunit of the cohesin complex and are required for cohesin-dependent insulation activity. Molecular and cellular biology 2011, 31: 2174–2183.

38. Saldaña-Meyer R, Rodriguez-Hernaez J, Escobar T, Nishana M, Jácome-López K, Nora EP, Bruneau BG, Tsirigos A, Furlan-Magaril M, Skok J: RNA Interactions Are Essential for CTCF-Mediated Genome Organization. Molecular cell 2019.

39. Cerami E, Gao J, Dogrusoz U, Gross BE, Sumer SO, Aksoy BA, Jacobsen A, Byrne CJ, Heuer ML, Larsson E: The cBio cancer genomics portal: an open platform for exploring multidimensional cancer genomics data. AACR; 2012.

40. Sati L, Zeiss C, Yekkala K, Demir R, McGrath J: Expression of the CTCFL Gene during Mouse Embryogenesis Causes Growth Retardation, Postnatal Lethality, and Dysregulation of the Transforming Growth Factor β Pathway. Molecular and cellular biology 2015, 35: 3436–3445.

41. Parelho V, Hadjur S, Spivakov M, Leleu M, Sauer S, Gregson HC, Jarmuz A, Canzonetta C, Webster Z, Nesterova T: Cohesins functionally associate with CTCF on mammalian chromosome arms. Cell 2008, 132: 422–433.

42. Rubio ED, Reiss DJ, Welcsh PL, Disteche CM, Filippova GN, Baliga NS, Aebersold R, Ranish JA, Krumm A: CTCF physically links cohesin to chromatin. Proceedings of the National Academy of Sciences 2008, 105: 8309–8314.

43. Wendt KS, Peters J-M: How cohesin and CTCF cooperate in regulating gene expression. Chromosome research 2009, 17: 201–214.

44. Kumar S, Srivastav RK, Wilkes DW, Ross T, Kim S, Kowalski J, Chatla S, Zhang Q, Nayak A, Guha M: Estrogen-dependent DLL1-mediated Notch signaling promotes luminal breast cancer. Oncogene 2019, 38: 2092.

45. Sales-Dias J, Silva G, Lamy M, Ferreira A, Barbas A: The Notch ligand DLL1 exerts carcinogenic features in human breast cancer cells. PloS one 2019, 14: e0217002.

46. Vasilaki E, Kanaki Z, Stravopodis DJ, Klinakis A: Dll1 Marks Cells of Origin of Ras-Induced Cancer in Mouse Squamous Epithelia. Translational oncology 2018, 11: 1213–1219.

47. Bettinsoli P, Ferrari-Toninelli G, Bonini S, Prandelli C, Memo M: Notch ligand Delta-like 1 as a novel molecular target in childhood neuroblastoma. BMC cancer 2017, 17: 352.

48. Xu H, Shan J, Jurukovski V, Yuan L, Li J, Tian K: TSP50 encodes a testis-specific protease and is negatively regulated by p53. Cancer research 2007, 67: 1239–1245.

49. Ramachandran A, Vizan P, Das D, Chakravarty P, Vogt J, Rogers KW, Müller P, Hinck AP, Sapkota GP, Hill CS: TGF-β uses a novel mode of receptor activation to phosphorylate SMAD1/5 and induce epithelial-to-mesenchymal transition. Elife 2018, 7: e31756.

50. Memon A, Lee WK: KLF10 as a Tumor Suppressor Gene and Its TGF-β Signaling. Cancers 2018, 10: 161.

51. Liu B, Sun X, Suyeoka G, Garcia JG, Leiderman YI: TGFβ signaling induces expression of Gadd45b in retinal ganglion cells. Investigative ophthalmology & visual science 2013, 54: 1061–1069.

52. Zhang Y, Liu Z: STAT1 in cancer: friend or foe? Discovery medicine 2017, 24: 19–29.

53. Plisov S, Tsang M, Shi G, Boyle S, Yoshino K, Dunwoodie SL, Dawid IB, Shioda T, Perantoni AO, de Caestecker MP: Cited1 is a bifunctional transcriptional cofactor that regulates early nephronic patterning. Journal of the American Society of Nephrology 2005, 16: 1632–1644.

54. Hansen AS, Amitai A, Cattoglio C, Tjian R, Darzacq X: Guided nuclear exploration increases CTCF target search efficiency. bioRxiv 2018: 495457.

55. Hansen AS, Hsieh T-HS, Cattoglio C, Pustova I, Saldaña-Meyer R, Reinberg D, Darzacq X, Tjian R: Distinct classes of chromatin loops revealed by deletion of an RNA-binding Region in CTCF. Molecular cell 2019.

56. Wutz G, Várnai C, Nagasaka K, Cisneros DA, Stocsits RR, Tang W, Schoenfelder S, Jessberger G, Muhar M, Hossain MJ: Topologically associating domains and chromatin loops depend on cohesin and are regulated by CTCF, WAPL, and PDS5 proteins. The EMBO journal 2017, 36: 3573–3599.

57. Garg R, Benedetti LG, Abera MB, Wang H, Abba M, Kazanietz MG: Protein kinase C and cancer: what we know and what we do not. Oncogene 2014, 33: 5225.

58. Bergmaier P, Weth O, Dienstbach S, Boettger T, Galjart N, Mernberger M, Bartkuhn M, Renkawitz R: Choice of binding sites for CTCFL compared to CTCF is driven by chromatin and by sequence preference. Nucleic acids research 2018, 46: 7097–7107.

59. Gong Y, Lazaris C, Sakellaropoulos T, Lozano A, Kambadur P, Ntziachristos P, Aifantis I, Tsirigos A: Stratification of TAD boundaries reveals preferential insulation of superenhancers by strong boundaries. Nature communications 2018, 9: 1–12.

60. Haarhuis JH, van der Weide RH, Blomen VA, Yáñez-Cuna JO, Amendola M, van Ruiten MS, Krijger PH, Teunissen H, Medema RH, van Steensel B: The cohesin release factor WAPL restricts chromatin loop extension. Cell 2017, 169: 693-707. e614.

61. Rao SS, Huang S-C, St Hilaire BG, Engreitz JM, Perez EM, Kieffer-Kwon K-R, Sanborn AL, Johnstone SE, Bascom GD, Bochkov ID: Cohesin loss eliminates all loop domains. Cell 2017, 171: 305-320. e324.

62. Pugacheva EM, Kubo N, Loukinov D, Tajmul M, Kang S, Kovalchuk AL, Strunnikov AV, Zentner GE, Ren B, Lobanenkov VV: CTCF mediates chromatin looping via N-terminal domain-dependent cohesin retention. Proceedings of the National Academy of Sciences 2020.

63. Zoabi M, Nadar-Ponniah PT, Khoury-Haddad H, Usaj M, Budowski-Tal I, Haran T, Henn A, Mandel-Gutfreund Y, Ayoub N: RNA-dependent chromatin localization of KDM4D lysine demethylase promotes H3K9me3 demethylation. Nucleic acids research 2014, 42: 13026–13038.

64. Schmidl C, Rendeiro AF, Sheffield NC, Bock C: ChIPmentation: fast, robust, low-input ChIP-seq for histones and transcription factors. Nature methods 2015, 12: 963.

65. Langmead B, Salzberg SL: Fast gapped-read alignment with Bowtie 2. Nat Methods 2012, 9: 357–359.

66. Zhang Y, Liu T, Meyer CA, Eeckhoute J, Johnson DS, Bernstein BE, Nusbaum C, Myers RM, Brown M, Li W: Model-based analysis of ChIP-Seq (MACS). Genome biology 2008, 9: R137.

67. Ramirez F, Ryan DP, Gruning B, Bhardwaj V, Kilpert F, Richter AS, Heyne S, Dundar F, Manke T: deepTools2: a next generation web server for deep-sequencing data analysis. Nucleic Acids Res 2016, 44: W160–165.

68. Stark R, Brown G: DiffBind: differential binding analysis of ChIP-Seq peak data. R package version 2011, 100: 4–3.

69. Dobin A, Davis CA, Schlesinger F, Drenkow J, Zaleski C, Jha S, Batut P, Chaisson M, Gingeras TR: STAR: ultrafast universal RNA-seq aligner. Bioinformatics 2013, 29: 15–21.

70. Love MI, Huber W, Anders S: Moderated estimation of fold change and dispersion for RNA-seq data with DESeq2. Genome biology 2014, 15: 550.

71. Larsson J: Eulerr: area-proportional Euler Diagrams. R Package version 4.1. 0. 2018.

72. Ruffier M, Kähäri A, Komorowska M, Keenan S, Laird M, Longden I, Proctor G, Searle S, Staines D, Taylor K: Ensembl core software resources: storage and programmatic access for DNA sequence and genome annotation. Database 2017, 2017.

73. Lazaris C, Kelly S, Ntziachristos P, Aifantis I, Tsirigos A: HiC-bench: comprehensive and reproducible Hi-C data analysis designed for parameter exploration and benchmarking. BMC genomics 2017, 18: 22.

74. Tsirigos A, Haiminen N, Bilal E, Utro F: GenomicTools: a computational platform for developing high-throughput analytics in genomics. Bioinformatics 2011, 28: 282–283.

75. Yu G, Wang L-G, He Q-Y: ChIPseeker: an R/Bioconductor package for ChIP peak annotation, comparison and visualization. Bioinformatics 2015, 31: 2382–2383.

76. Bailey TL, Boden M, Buske FA, Frith M, Grant CE, Clementi L, Ren J, Li WW, Noble WS: MEME SUITE: tools for motif discovery and searching. Nucleic acids research 2009, 37: W202–W208.

77. Machanick P, Bailey TL: MEME-ChIP: motif analysis of large DNA datasets. Bioinformatics 2011, 27: 1696–1697.

78. Heinz S, Benner C, Spann N, Bertolino E, Lin YC, Laslo P, Cheng JX, Murre C, Singh H, Glass CK: Simple combinations of lineage-determining transcription factors prime cisregulatory elements required for macrophage and B cell identities. Molecular cell 2010, 38: 576–589.

79. Imakaev M, Fudenberg G, McCord RP, Naumova N, Goloborodko A, Lajoie BR, Dekker J, Mirny LA: Iterative correction of Hi-C data reveals hallmarks of chromosome organization. Nature methods 2012, 9: 999.

80. Crane E, Bian Q, McCord RP, Lajoie BR, Wheeler BS, Ralston EJ, Uzawa S, Dekker J, Meyer BJ: Condensin-driven remodelling of X chromosome topology during dosage compensation. Nature 2015, 523: 240.

81. Durand NC, Robinson JT, Shamim MS, Machol I, Mesirov JP, Lander ES, Aiden EL: Juicebox provides a visualization system for Hi-C contact maps with unlimited zoom. Cell systems 2016, 3: 99–101.

82. Durand NC, Shamim MS, Machol I, Rao SS, Huntley MH, Lander ES, Aiden EL: Juicer provides a one-click system for analyzing loop-resolution Hi-C experiments. Cell systems 2016, 3: 95–98.

83. Klenova EM, Nicolas RH, Sally U, Carne AF, Lee RE, Lobanenkov VV, Goodwin GH: Molecular weight abnormalities of the CTCF transcription factor: CTCF migrates aberrantly in SDS-PAGE and the size of the expressed protein is affected by the UTRs and sequences within the coding region of the CTCF gene. Nucleic acids research 1997, 25: 466–473.

84. Skok J, Nishana M, Rodriguez-Hernaez J. The zinc finger and N/C terminal domains of CTCF and CTCFL contribute to binding site-specific functional impact. GSE140363, Gene Expression Omnibus. 2019; Available from: https://www.ncbi.nlm.nih.gov/geo/query/acc.cgi?acc=GSE140363

