## Supplemental Figures for "Defining the relative and combined contribution of CTCF and CTCFL to genomic regulation"

Fig S1

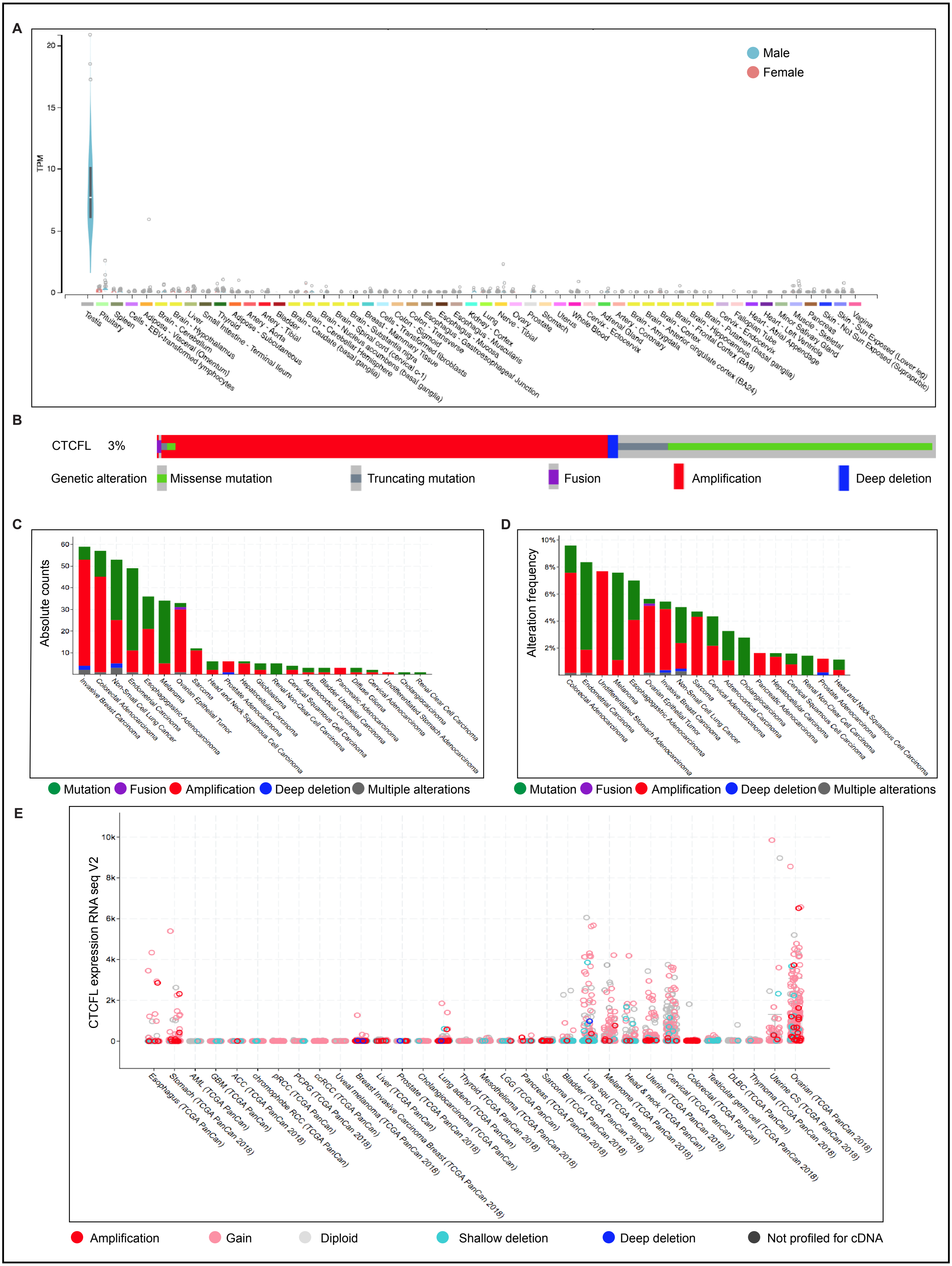
A
TPM
Male
Female
Testis
Pituitary
Spleen
Cells - EBV-transformed lymphocytes
Adipose - Visceral (Omentum)
Brain - Cerebellum
Brain - Hypothalamus
Liver
Small Intestine - Terminal Ileum
Thyroid
Adipose - Subcutaneous
Artery - Aorta
Artery - Tibial
Bladder
Brain - Caudate (basal ganglia)
Brain - Cerebellar Hemisphere
Brain - Nucleus accumbens (basal ganglia)
Brain - Spinal cord (cervical c-1)
Brain - Substantia nigra
Breast - Mammary Tissue
Cells - Transformed fibroblasts
Colon - Sigmoid
Colon - Transverse
Esophagus - Gastroesophageal Junction
Esophagus - Mucosa
Esophagus - Muscularis
Kidney - Cortex
Lung
Nerve - Tibial
Ovary
Prostate
Stomach
Uterus
Whole Blood
Cervix - Ectocervix
Adrenal Gland
Artery - Coronary
Brain - Amygdala
Brain - Anterior cingulate cortex (BA24)
Brain - Cortex
Brain - Frontal Cortex (BA9)
Brain - Hippocampus
Brain - Putamen (basal ganglia)
Cervix - Endocervix
Fallopian Tube
Heart - Atrial Appendage
Heart - Left Ventricle
Minor Salivary Gland
Muscle - Skeletal
Pancreas
Skin - Not Sun Exposed (Suprapubic)
Skin - Sun Exposed (Lower leg)
Vagina
B
CTCFL 3%
Genetic alteration
Missense mutation
Truncating mutation
Fusion
Amplification
Deep deletion
C
Absolute counts
Invasive Breast Carcinoma
Colorectal Adenocarcinoma
Non-Small Cell Lung Cancer
Endometrial Carcinoma
Esophagogastric Adenocarcinoma
Melanoma
Ovarian Epithelial Tumor
Sarcoma
Head and Neck Squamous Cell Carcinoma
Prostate Adenocarcinoma
Hepatobiliary Cancer
Glioblastoma
Renal Non-Clear Cell Carcinoma
Cervical Squamous Cell Carcinoma
Adrenocortical Carcinoma
Bladder Urothelial Carcinoma
Pancreatic Adenocarcinoma
Diffuse Glioma
Cervical Adenocarcinoma
Undifferentiated Stomach Adenocarcinoma
Cholangiocarcinoma
Renal Clear Cell Carcinoma
Mutation
Fusion
Amplification
Deep deletion
Multiple alterations
D
Alteration frequency
Colorectal Adenocarcinoma
Endometrial Carcinoma
Undifferentiated Stomach Adenocarcinoma
Melanoma
Esophagogastric Adenocarcinoma
Ovarian Epithelial Tumor
Invasive Breast Carcinoma
Non-Small Cell Lung Cancer
Sarcoma
Cervical Adenocarcinoma
Adrenocortical Carcinoma
Cholangiocarcinoma
Pancreatic Adenocarcinoma
Hepatocellular Carcinoma
Cervical Squamous Cell Carcinoma
Renal Non-Clear Cell Carcinoma
Prostate Adenocarcinoma
Head and Neck Squamous Cell Carcinoma
Mutation
Fusion
Amplification
Deep deletion
Multiple alterations
E
CTCFL expression RNA seq V2
Esophagus (TCGA PanCan)
Stomach (TCGA PanCan 2018)
AML (TCGA PanCan)
GBM (TCGA PanCan)
ACC (TCGA PanCan 2018)
chromophobe RCC (TCGA PanCan)
pRCC (TCGA PanCan)
PCPG (TCGA PanCan 2018)
ccRCC (TCGA PanCan)
Uveal melanoma (TCGA PanCan 2018)
Breast Invasive Carcinoma Breast (TCGA PanCan 2018)
Liver (TCGA PanCan)
Prostate (TCGA PanCan 2018)
Cholangiocarcinoma (TCGA PanCan)
Lung adeno (TCGA PanCan 2018)
Thyroid (TCGA PanCan)
Mesothelioma (TCGA PanCan 2018)
LGG (TCGA PanCan)
Pancreas (TCGA PanCan 2018)
Sarcoma (TCGA PanCan 2018)
Bladder (TCGA PanCan 2018)
Lung squ (TCGA PanCan)
Melanoma (TCGA PanCan 2018)
Head & neck (TCGA PanCan)
Uterine (TCGA PanCan 2018)
Cervical (TCGA PanCan)
Colorectal (TCGA PanCan)
Testicular germ cell (TCGA PanCan 2018)
DLBC (TCGA PanCan)
Thymoma (TCGA PanCan 2018)
Uterine CS (TCGA PanCan 2018)
Ovarian (TCGA PanCan 2018)
Amplification
Gain
Diploid
Shallow deletion
Deep deletion
Not profiled for cDNA

Fig S2

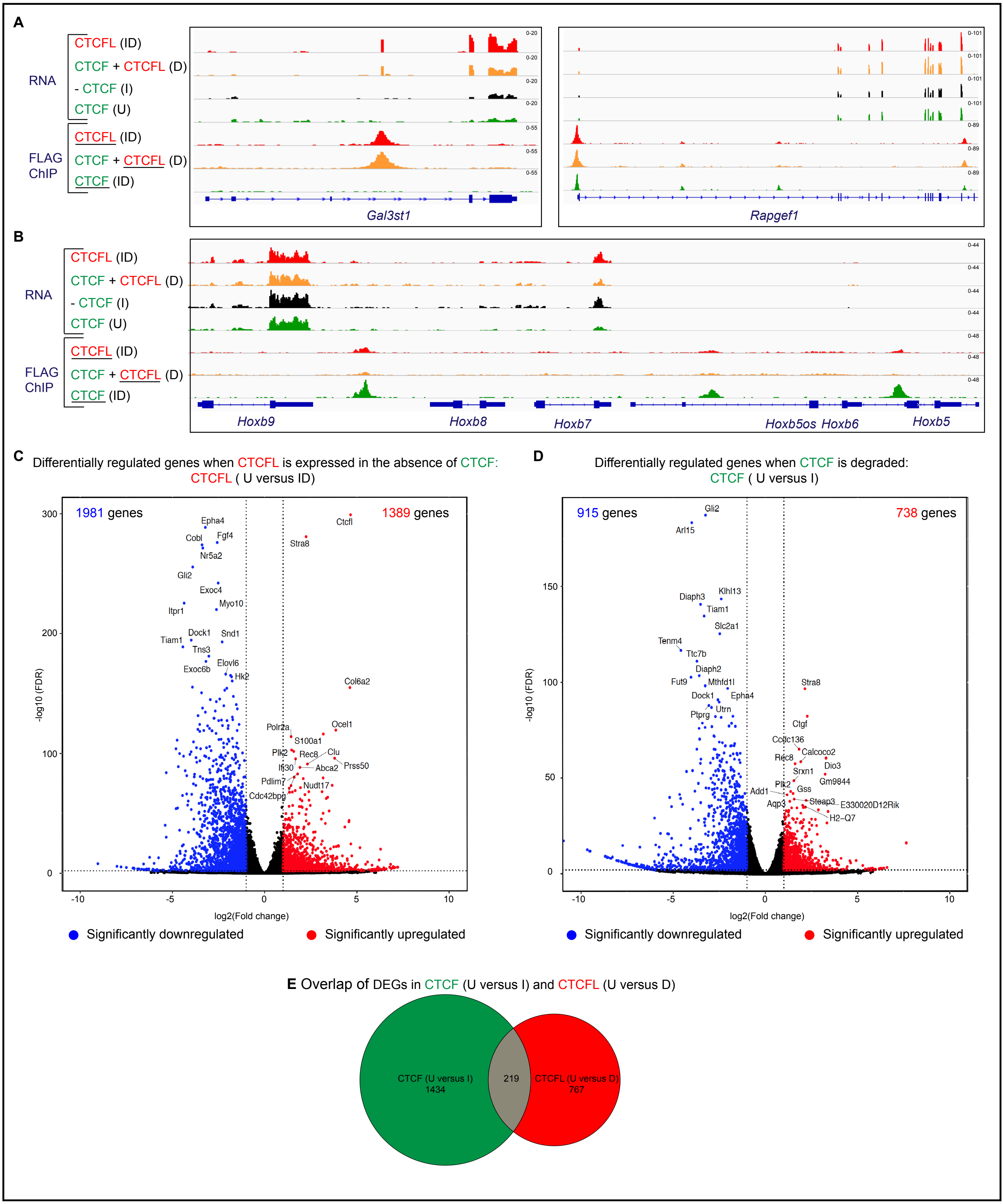
A
RNA
CTCFL (ID)
CTCF + CTCFL (D)
- CTCF (I)
CTCF (U)
FLAG ChIP
CTCFL (ID)
CTCF + CTCFL (D)
CTCF (ID)
Gal3st1
Rapgef1
B
Hoxb9
Hoxb8
Hoxb7
Hoxb5os Hoxb6
Hoxb5
C Differentially regulated genes when CTCFL is expressed in the absence of CTCF:
CTCFL ( U versus ID)
1981 genes
1389 genes
-log10 (FDR)
log2(Fold change)
D Differentially regulated genes when CTCF is degraded:
CTCF ( U versus I)
915 genes
738 genes
-log10 (FDR)
log2(Fold change)
Significantly downregulated
Significantly upregulated
E Overlap of DEGs in CTCF (U versus I) and CTCFL (U versus D)
CTCF (U versus I)
1434
219
CTCFL (U versus D)
767

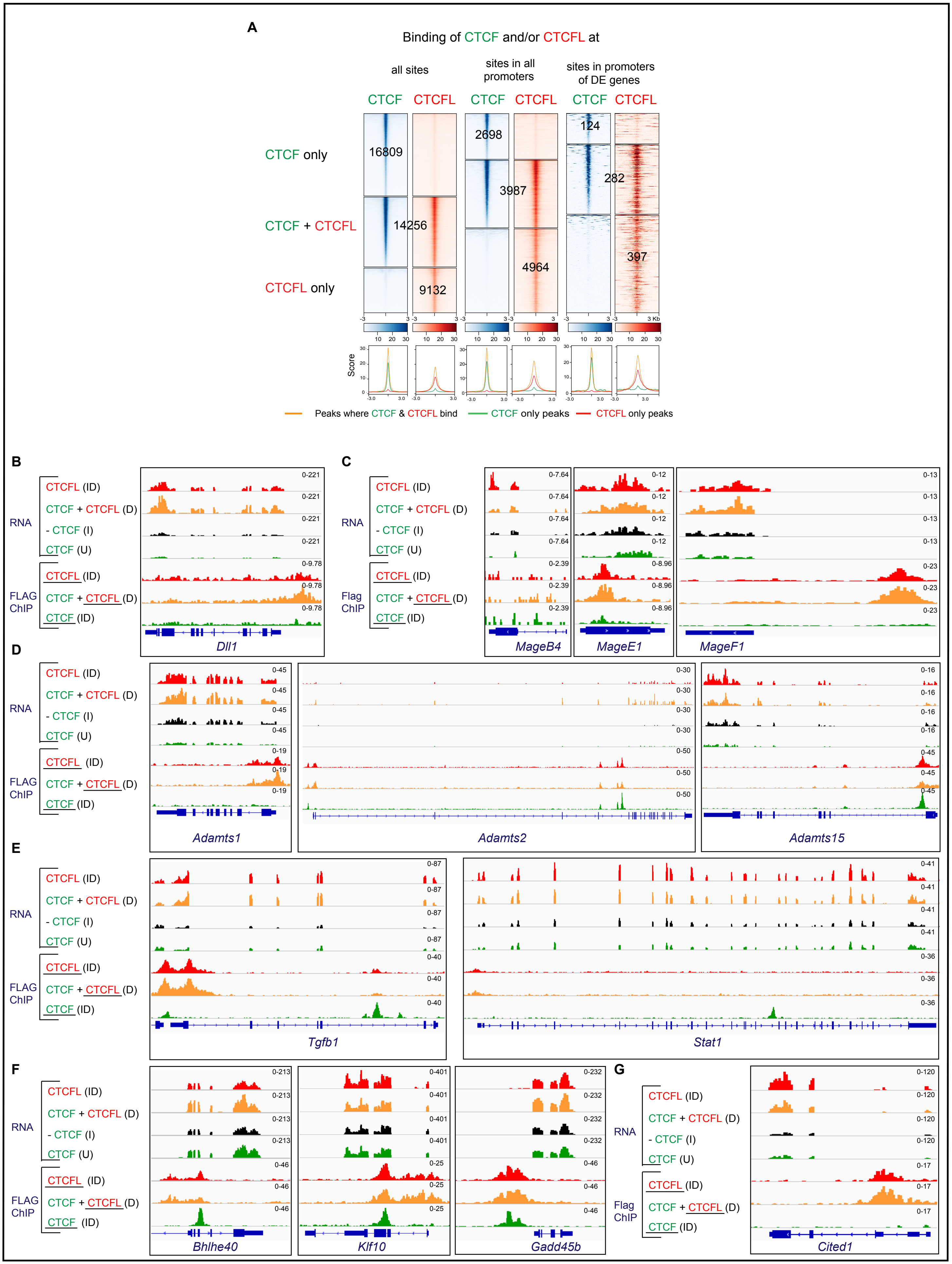

A
Binding of CTCF and/or CTCFL at
all sites
sites in all promoters
sites in promoters of DE genes
CTCF only
CTCF + CTCFL
CTCFL only
16809
14256
9132
2698
3987
4964
124
282
397
Score
Peaks where CTCF & CTCFL bind
CTCF only peaks
CTCFL only peaks
B
C
D
E
F
G
RNA
FLAG ChIP
Flag ChIP
CTCFL (ID)
CTCF + CTCFL (D)
- CTCF (I)
CTCF (U)
CTCF (ID)
Dll1
MageB4
MageE1
MageF1
Adamts1
Adamts2
Adamts15
Tgfb1
Stat1
Bhlhe40
Klf10
Gadd45b
Cited1

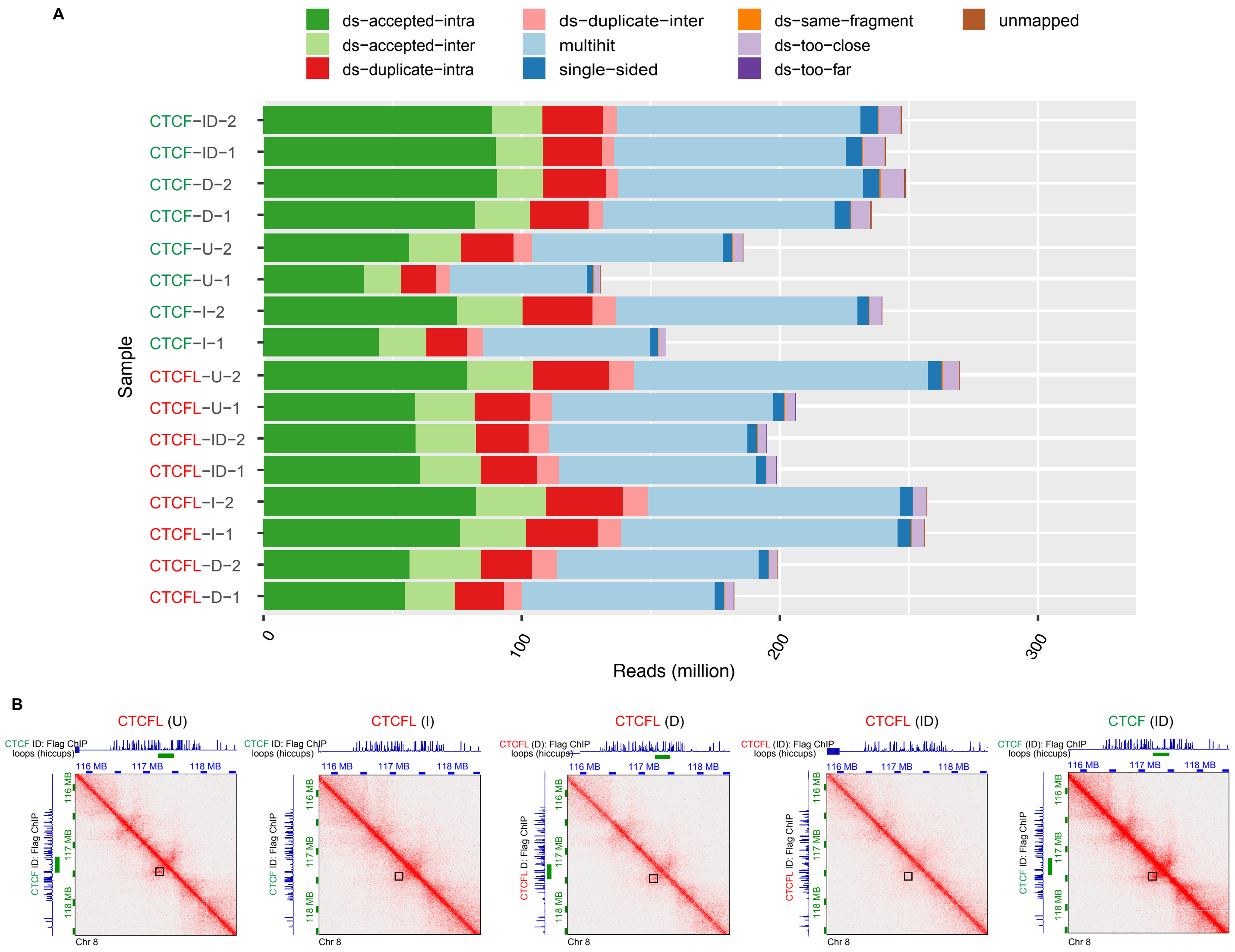
A
ds−accepted−intra
ds−accepted−inter
ds−duplicate−intra
ds−duplicate−inter
multihit
single−sided
ds−same−fragment
ds−too−close
ds−too−far
unmapped
Sample
CTCF−ID−2
CTCF−ID−1
CTCF−D−2
CTCF−D−1
CTCF−U−2
CTCF−U−1
CTCF−I−2
CTCF−I−1
CTCFL−U−2
CTCFL−U−1
CTCFL−ID−2
CTCFL−ID−1
CTCFL−I−2
CTCFL−I−1
CTCFL−D−2
CTCFL−D−1
0
100
200
300
Reads (million)
B
CTCFL (U)
CTCFL (I)
CTCFL (D)
CTCFL (ID)
CTCF (ID)
CTCF ID: Flag ChIP
loops (hiccups)
CTCFL (D): Flag ChIP
CTCFL (ID): Flag ChIP
CTCF (ID): Flag ChIP
116 MB
117 MB
118 MB
Chr 8

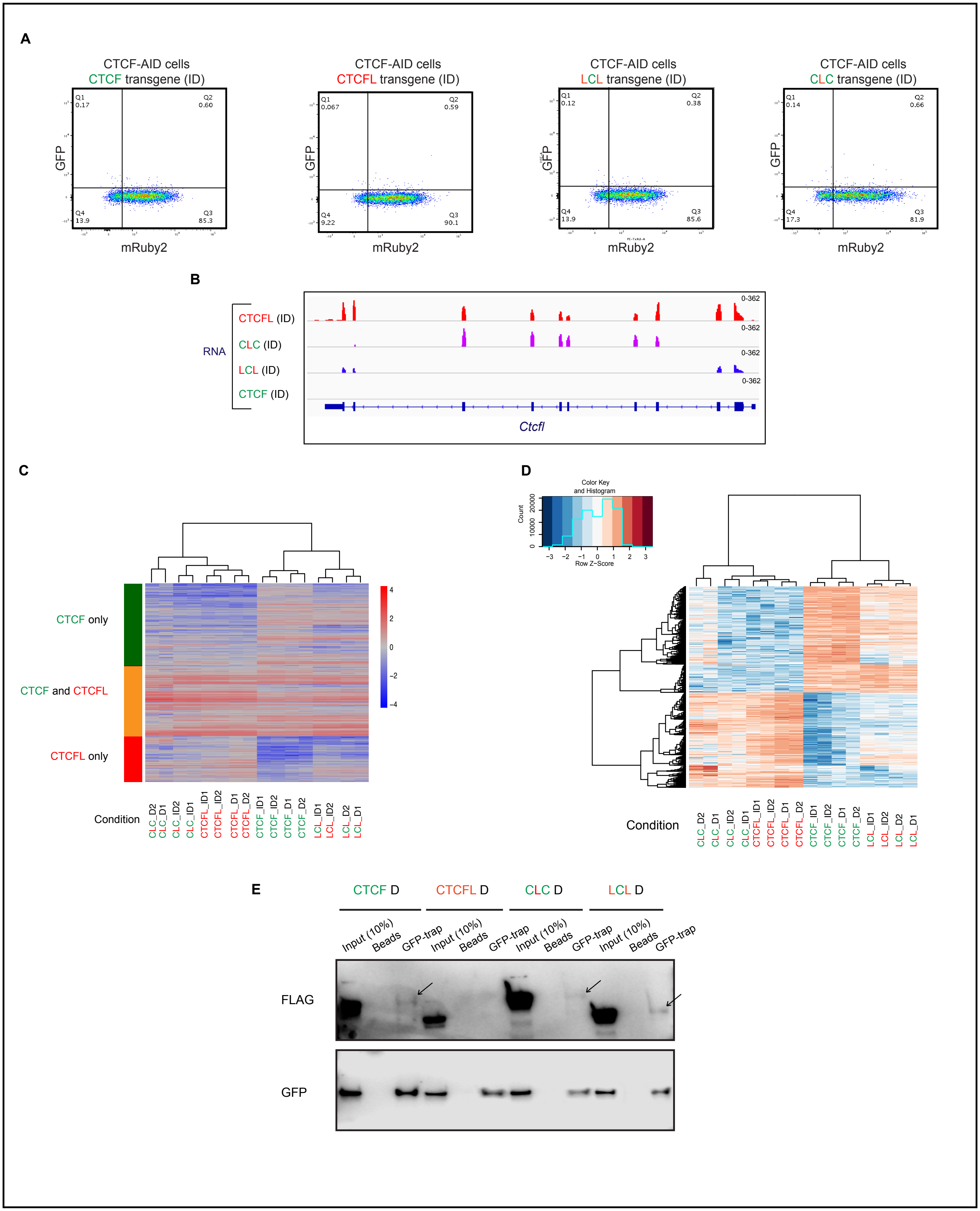
A
CTCF-AID cells
CTCF transgene (ID)
CTCFL transgene (ID)
LCL transgene (ID)
CLC transgene (ID)
GFP
mRuby2
Q1
Q2
Q3
Q4
0.17
0.60
13.9
85.3
0.067
0.59
9.22
90.1
0.12
0.38
13.9
85.6
0.14
0.66
17.3
81.9
B
RNA
CTCFL (ID)
CLC (ID)
LCL (ID)
CTCF (ID)
0-362
Ctcfl
C
CTCF only
CTCF and CTCFL
CTCFL only
Condition
CLC_D2
CLC_D1
CLC_ID2
CLC_ID1
CTCFL_ID1
CTCFL_ID2
CTCFL_D1
CTCFL_D2
CTCF_ID1
CTCF_ID2
CTCF_D1
CTCF_D2
LCL_ID1
LCL_ID2
LCL_D2
LCL_D1
D
Color Key and Histogram
Count
Row Z-Score
Condition
E
CTCF D
CTCFL D
CLC D
LCL D
Input (10%)
Beads
GFP-trap
FLAG
GFP

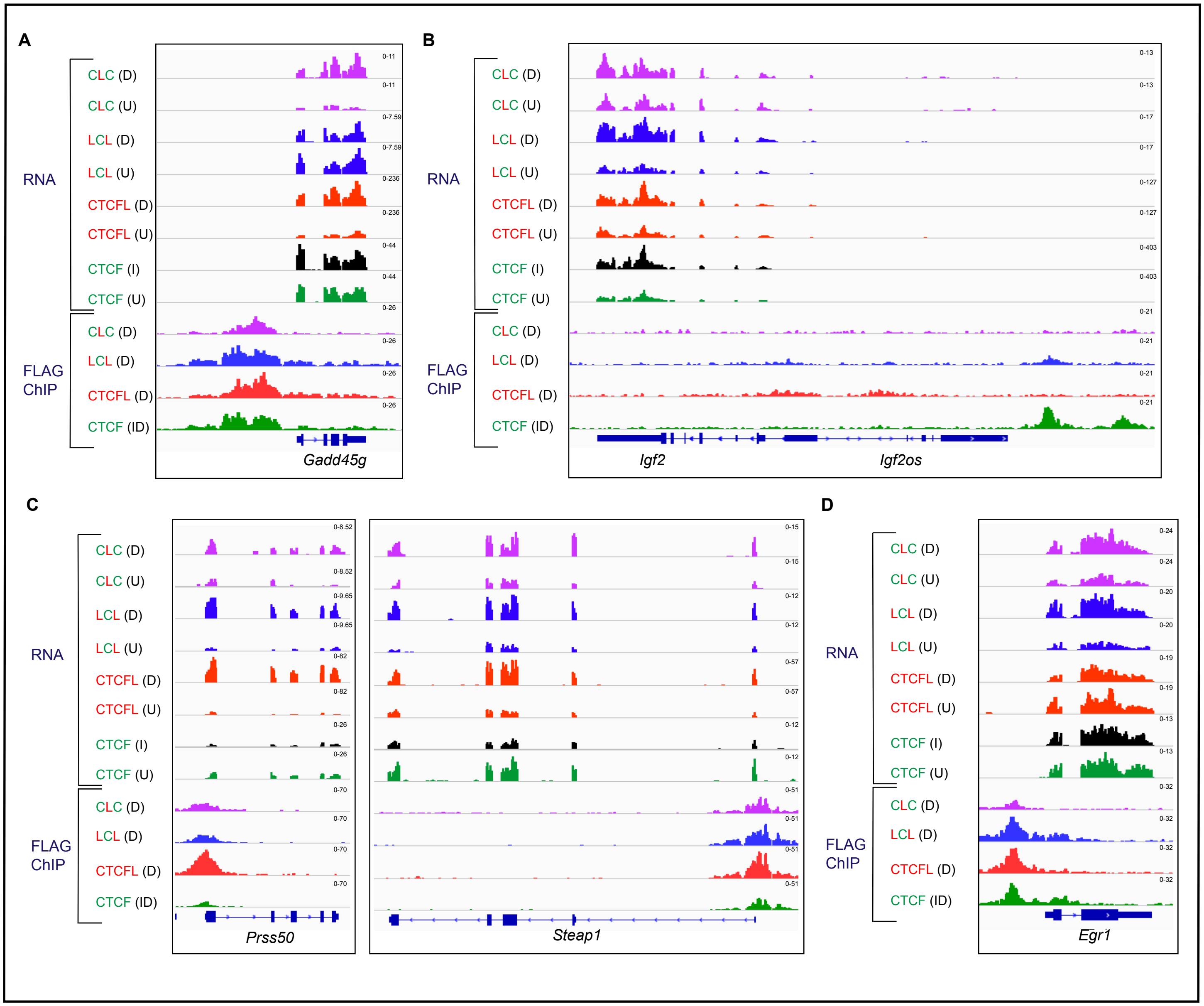
A
RNA
CLC (D)
CLC (U)
LCL (D)
LCL (U)
CTCFL (D)
CTCFL (U)
CTCF (I)
CTCF (U)
FLAG ChIP
CLC (D)
LCL (D)
CTCFL (D)
CTCF (ID)
Gadd45g
B
Igf2
Igf2os
C
Prss50
Steap1
D
Egr1

Fig S7

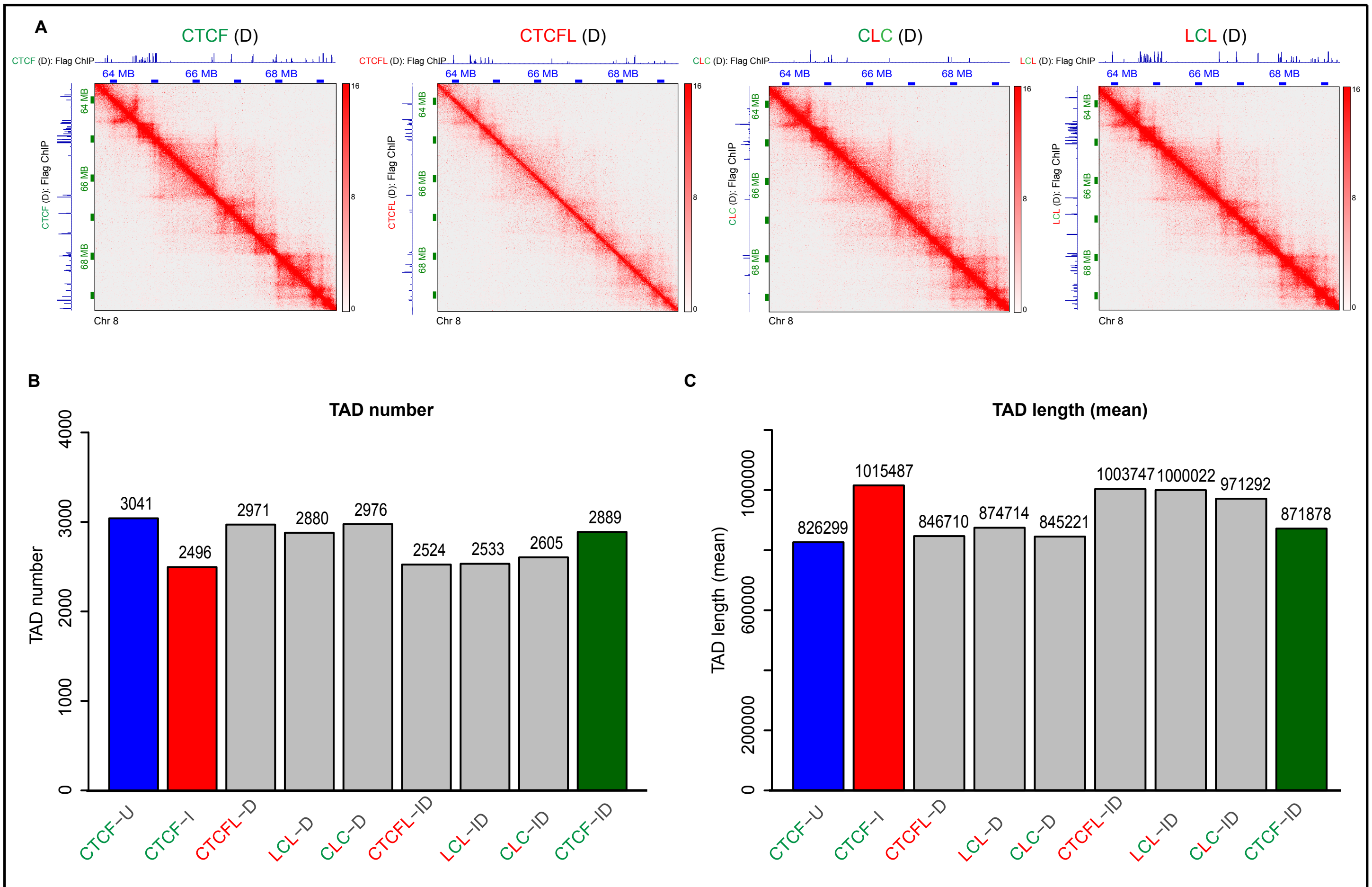
A
CTCF (D)
CTCFL (D)
CLC (D)
LCL (D)
CTCF (D): Flag ChIP
CTCFL (D): Flag ChIP
CLC (D): Flag ChIP
LCL (D): Flag ChIP
64 MB
66 MB
68 MB
Chr 8
16
8
0
B
TAD number
TAD number
0
1000
2000
3000
4000
3041
2496
2971
2880
2976
2524
2533
2605
2889
CTCF−U
CTCF−I
CTCFL−D
LCL−D
CLC−D
CTCFL−ID
LCL−ID
CLC−ID
CTCF−ID
C
TAD length (mean)
TAD length (mean)
0
200000
600000
1000000
826299
1015487
846710
874714
845221
1003747
1000022
971292
871878
CTCF−U
CTCF−I
CTCFL−D
LCL−D
CLC−D
CTCFL−ID
LCL−ID
CLC−ID
CTCF−ID

### Supplementary Figure Legends

**Fig S1. CTCFL is genetically altered and aberrantly expressed in numerous cancers. A** CTCFL is expressed only in testes in healthy individuals. Gene expression data for CTCFL (ENSG00000124092.8) was retrieved from GtexPortal (GTEx Analysis Release V7; dbGaP Accession phs000424.v7.p2). Data is sorted by median expression on a linear scale with males marked by blue and females by pink. **B** *CTCFL* is altered in 3% of cancer patients/samples profiled in cBioportal. Data was retrieved from Oncoprint of cBioPortal. The distribution of various alterations (missense mutation, truncations, fusion, amplification and deletion) in *CTCFL* in cancer are shown. As of 06/30/19, 3% of cancer patients profiled in cBioPortal had an alteration in *CTCFL*, the ratio of altered / profiled being 382 / 10950. **C, D** *CTCFL* is amplified across several cancer types. Alterations such as mutation, fusion, amplification and deletion reported in cancer patients are color coded. Absolute count of the patients (**C**) and alteration frequency in percentage (**D**) in various cancer types are shown. **E** CTCFL expression (RNA seq) and copy number variation in several cancer types. Data is sorted by median on a linear scale. Copy number alterations such as amplification, gain, diploid, deep and shallow deletion etc. are color coded.

**Fig S2. CTCF and CTCFL bind to and regulate distinct sets of genes. A, B** IgV tracks showing RNA-seq in all four conditions (U, I, D and ID) in cells harboring the CTCFL transgene. ChIP-seq using FLAG antibody to detect the binding of transgenic CTCFL and CTCF. Binding and expression at *Gal3st1* and *Rapgef1* (**A**) and the *Hoxb* cluster (**B**) is shown. **C, D** Volcano plots showing differentially expressed genes in the wild-type (U) versus CTCFL expressing mESCs in the absence of endogenous CTCF (ID condition) (**C**) and wild type (U) versus CTCF degraded (I condition) cells (**D**). Red and blue points mark the genes with significantly increased or decreased expression respectively (FDR<0.01). The x-axis shows the log2 fold-change in expression and the y-axis the log

10 (False discovery rate) of a gene being differentially expressed. The number of genes that are significantly up or downregulated are indicated in either case. **E** Venn diagrams showing the overlap of differentially expressed genes (DEG) between U versus I condition and U versus D condition (in cells harboring the CTCFL transgene).

**Fig S3. CTCFL activates cancer testes antigens (CTA) and components of cancer relevant signaling pathways. A** Distribution of peaks bound exclusively by CTCF and/or CTCFL at all binding sites, binding sites in promoters and in promoters of differentially expressed genes (DEG) when CTCFL is expressed (D condition). Average profiles are shown below the corresponding heatmaps. **B-G** IgV tracks showing RNA-seq in all four conditions (U, I, D and ID) in cells harboring the CTCFL transgene. ChIP-seq was performed using a FLAG antibody to detect the binding of transgenic CTCFL and CTCF. Binding of CTCFL and CTCF and transcriptional changes are shown for *Dll1* (**B**) *MageB4, MageE1, MageF1* (**C**) *Adamts1, Adamts2, Adamts15* (**D**) *Tgfb1* and *Stat1* (**E**) *Bhlhe40, Klf10* and *Gadd45b* (**F**) and *Cited1* (**G**).

**Fig S4. The impact of CTCFL on 3D chromatin organization. A** Hi-C analysis was performed in cells harboring CTCF and CTCFL transgenes under U, I, D and ID conditions in replicates. Statistical report showing the number (in millions) of reads assigned to the different categories following the filtering of reads using HiC bench [73]. Only “ds accepted-intra” and “ds accepted inter” reads were used for further analysis. **B** Snapshot of HiC data from Juicebox showing loss of loops when CTCF is degraded (CTCFL-I) and rescue under conditions of CTCFL-D and CTCF-ID, but not CTCFL-ID. The loop regions from Hiccups and Flag ChIPs corresponding to the respective cell lines are shown along the x-and y-axis.

**Fig S5. The role of CTCF and CTCFL zinc fingers and N/C terminal regions in site specific binding. A** Flow cytometry confirms the comparable level of mRuby2 expression of transgenic CTCF,

CTCFL, LCL and CLC. **B** IgV tracks of RNA-seq showing expression of the transgenes. At the *Ctcfl* locus, peaks are seen at the exons corresponding to the swapped regions with the chimeric proteins, all exons with *Ctcfl* and no peaks with the *Ctcf* transgene. **C, D** Heatmap showing the clustering of ChIP-seq signals where CTCF, CTCFL, CLC and LCL bind in the presence (D) and absence (ID) of endogenous CTCF. The peak sites corresponding to CTCF only, CTCFL only and CTCF and CTCFL overlapping sites were used for the analysis. These were sorted by site in (**C**) but not in (**D**). **E**. Immunoprecipitation of endogenous GFP tagged CTCF using GFP-Trap and blotting with FLAG antibody demonstrate an interaction between CTCF and transgenic FLAG tagged CTCF, CLC and LCL but not CTCFL. The blot with the GFP antibody shows the pull down of endogenous CTCF in cells expressing the different transgenes.

**Fig S6. Impact of fusion proteins on gene expression. A-D** IgV tracks showing RNA-seq of indicated conditions in cells harboring CTCF, CTCFL, CLC or LCL transgene under the indicated conditions. ChIP-seq using FLAG antibody was used to detect the binding of transgenes. Binding and expression at *Gadd45g* (**A**), *Igf2* (**B**), *Prss50*, *Steap1* (**C**) and *Egr1* (**D**) is shown.

**Fig S7. The impact of fusion proteins on chromatin organization. A** Snapshot of Hi-C data from Juicebox corresponding to Chr 8: 63,616,214-69,456,200 8: 63,566,214- 69,406,200 at 10 kb resolution. Cells harboring CTCF, CTCFL, CLC or LCL transgenes were treated with Dox as indicated for 4 days. The corresponding FLAG ChIPs are shown on the x-and y-axis as indicated. **B** The total number (**B**) and mean length (**C**) of TADs called in cells harboring transgenic CTCF, CTCFL, CLC and LCL when treated with D or ID as indicated for 4 days. Untreated (U) as well as CTCF degraded (I) cells harboring CTCF transgenes serve as positive and negative controls.
